# Oligodendrocytes support functional development of subcortical premotor neurons and navigation

**DOI:** 10.1101/2025.11.01.686017

**Authors:** Franziska Auer, Yifei Zhang, David Schoppik

## Abstract

Oligodendrocytes and the myelin they produce are required for proper balance behaviors in adult vertebrates. If, when, and how they shape the functional development of vestibular (balance) circuits and associated behaviors is unclear. We first adopted a pharmacological approach to investigate the contribution of oligodendrocytes across early development of larval zebrafish (*Danio rerio*). Disruption of oligodendrogenesis reduces the number of oligodendrocytes near vestibular neurons across development, but only impacts vertical navigation — a behavior requiring vestibular function — in older larvae. Correspondingly, we found that midbrain premotor neurons in the vertical navigation circuit only decreased their responses to tilts at later ages. In contrast, we found that the vestibulo-ocular reflex was unaffected by drug treatment, and that body-tilt responses in the underlying hindbrain vestibular neurons and their target extraocular motor neurons were unchanged. Targeted photoablation of oligodendrocytes in the anterior hindbrain replicated our pharmacological findings, selectively disrupting vertical navigation in older animals. By dissociating where and when developmentally-disrupted oligodendrogenesis impacts neuronal function and behavior, our work takes a major step towards understanding how oligodendrocytes enable the maturation of sensorimotor behaviors.

## INTRODUCTION

Oligodendrocytes — the myelin-producing cells of the central nervous system — ensheath axons of subcortical neurons responsible for the vital reflexes that control posture and gaze stabilization^1^. Myelin loss is profoundly disruptive to these vestibular (balance) behaviors^2–4^. Across early development, myelination and improved postural stability are concurrent^5,6^, suggesting that oligodendrocytes are foundational for vestibular circuit maturation. The complexity and *in utero* development of most vertebrate models hinder early investigation of brainstem function and vestibular behaviors. Consequentially, we do not know if, where, and how subcortical oligodendrocytes contribute to vestibular circuit development and associated behavioral maturation^7^.

Vestibular reflexes are shaped by lifelong sensorimotor learning^8,9^. Recent studies of adult sensory and motor learning offer models for how oligodendrocytes impact vestibular circuit maturation and behavior. Oligodendrogenesis might impact the representation of sensed body tilts as sensory deprivation reshapes myelin^10^ and enrichment increases myelination^11^. Complementarily, motor learning increases myelination^12,13^, oligodendrogenesis is key to skill acquisition^14–16^, and comparable findings hold for other forms of learning^17,18^. Oligodendrogenesis might therefore influence the premotor components of developing vestibular behaviors.

Alternatively, the earliest contributions of oligodendrocytes might be different than in mature animals^19,20^. Early oligodendrogenesis and myelination are sophisticated and regulated processes capable of shaping functional maturation of balance circuits and associated behavior^21–25^. Further, studies of monocular deprivation show that disrupting myelin-derived signaling aberrantly extends the critical period for visual development^26–28^ and destabilizes tuning, i.e. the set of sensory stimuli to which neurons respond^29^. Blocking adolescent oligodendrogenesis in rodents similarly extends the critical period in visual cortex^30^; comparable findings obtain in other cortical areas^31,32^. If subcortical oligodendrocytes behave similarly, then oligodendrogenesis could improve behavior by stabilizing the responses of developing vestibular neurons to body tilts.

Vertebrates, including the small model zebrafish (*Danio rerio*), use evolutionarily ancient subcortical vestibular circuits to balance^33^. In parallel, zebrafish oligodendrocytes use conserved mechanisms^34^ to progressively myelinate these circuits^35^. Over the first weeks of life, larvae progressively gain the ability to maintain posture^36–40^, coordinate their fins & trunk^41,42^, navigate up/down in the water^43^, and stabilize gaze^44–46^. These behaviors all rely on conserved vestibular circuits^47,48^ to sense, relay, and transform body tilts into both volitional and reflexive locomotion.

The subcortical circuits that control the gaze-stabilizing vestibulo-ocular reflex and vertical navigation overlap. Specifically, individual hindbrain interneurons in the tangential vestibular nucleus (TAN) target two areas^45,49–51^. First, they project to extraocular motor neurons in cranial nuclei nIII/nIV that counter-rotate the eyes to stabilize gaze after body tilts^46^. Additionally, they project to descending premotor neurons in the nucleus of the medial longitudinal fasciculus^52^, also known as the interstitial nucleus of Cajal (nMLF/INC)^53^. Loss of neurons in both the TAN and the nMLF/INC disrupts postural stability^39^ and vertical navigation^43^. Similarly, neurons in the TAN are indispensable for the vestibuloocular reflex^45^ and activation of the TAN leads to eye rotations^49^. Dissociable links between circuit architecture and associated behaviors offer functional and behavioral approaches to localize the consequences of disrupted oligodendrogenesis in both space and developmental time.

We adopted a pharmacological approach to disrupt oligodendrogenesis and measured the behavioral and neuronal consequences as larvae began to swim (4–10 days post fertilization). Treatment with the 2-oxoglutarate oxygenase inhibitor IOX1^54^ decreased the number of oligodendrocytes near balance neuron axons. Initially, vertical navigation in IOX1-treated fish was normal; with time, the impact of treatment became progressively more pronounced. Similarly, the neurons in the nMLF/INC of IOX1-treated fish decreased their response to nose-up tilts, but only at later ages. In contrast, responses of neurons in the TAN, their target neurons in nIII/nIV, and vestibulo-ocular reflex performance were all robust to IOX1 treatment. Finally, targeted photoablation of oligodendrocytes in the anterior medial longitudinal fasciculus (MLF) replicated our pharmacological finding, as they disrupted vertical navigation, but only in older animals. Together, our data reveals where and when loss of oligodendrocytes does and doesn’t impact maturation of vestibular neuron responses, vertical navigation, and the vestibulo-ocular reflex. This work significantly advances our understanding of how developing oligodendrocytes shape the maturation of sensorimotor circuits and associated behaviors.

## RESULTS

### IOX1 treatment disrupts oligodendrogenesis in balance-associated regions of the midbrain and hindbrain

We adopted a pharmacological approach to disrupt oligodendrogenesis that built on previous work showing that IOX1 reduces oligodendrocyte numbers^54^ without disrupting axonal branch patterning or synaptic deficits^55^. We exposed larvae to 4 µM IOX1 in 0.2% DMSO (drug) or DMSO only (control) starting at 54 hours post-fertilization (hpf), just before the first oligodendrocytes appear^56^. At 7 days post-fertilization (dpf), continuous IOX1 exposure reduced the number of oligodendrocytes (Figures 1A to 1C). We saw no effects on oligodendrocytes with later, discontinuous, or lower doses of IOX1 (Figures S1A to S1E).

**Figure 1:**
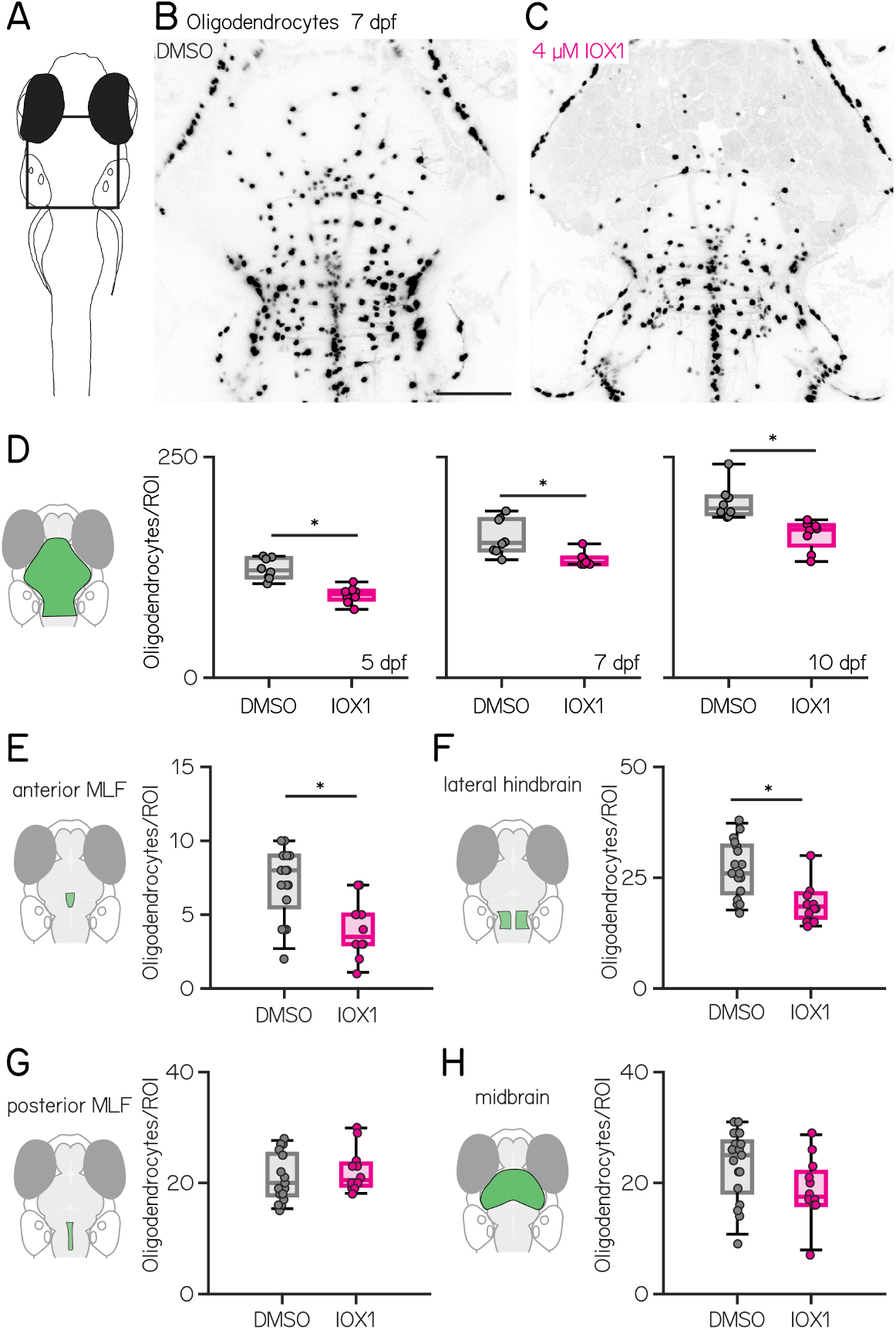
IOX1 treatment disrupts oligodendrogenesis in balance-associated regions of the midbrain and hindbrain. **(A)** Schematic of a larval zebrafish, viewed from above. Rectangle denotes the region in B&C. **(B–C)** Confocal images of 7 days post-fertilization (dpf) larval zebrafish with a nuclear-localized fluorophore expressed in oligodendrocytes. B (control) was treated with 0.2% DMSO and C (drug) was treated with 4 µM IOX1 in 0.2% DMSO. Scale bar 100 µm **D** Oligodendrocyte counts in 5, 7, and 10 dpf control (gray) and 4 µM IOX1-treated larvae (pink) (DMSO vs IOX1: 5 dpf: 122 [114 – 136] vs. 95 [89 – 99], p-value: 0.0005; 7 dpf: 153 [145 – 180] vs. 131 [129 – 137], p-value: 0.0033; 10 dpf: 192 [186 – 206] vs. 168 [150 – 173] oligodendrocytes, p-value: 0.0005) **(E-H)** Oligodendrocyte counts at 7 dpf, green shaded areas indicating the region of interest for: **(E)** anterior MLF (DMSO vs. IOX1: 8 [6 – 9] vs. 4 [3 – 5] p-value: 0.0077). **(F)** lateral hindbrain (DMSO vs. IOX1: 26 [22 – 33] vs. 19 [16 – 22]; p-value: 0.0084). **(G)** posterior MLF (DMSO vs. IOX1: 20 [18 – 25] vs. 21 [20 – 24]; p-value: 0.4110). **(H)**midbrain (DMSO vs. IOX1: 25 [18 – 28] vs. 18 [16 – 22]; p-value: 0.1221). * indicates p-value < 0.05.

Central balance circuits can be circumscribed by a region of interest that includes nuclei in the midbrain and hindbrain. To quantify the impact of IOX1 exposure on balance circuits we manually counted somata in this region at 5, 7, and 10 dpf using larvae that express a nuclear-localized fluorescent reporter in oligodendrocytes. At all time points, IOX1 treatment reduced the number of oligodendrocytes (Figure 1D; DMSO vs IOX1: 5 dpf: n = 8/8 DMSO/IOX1-treated animals; data shown as median [Q1 – Q3], 122 [114 – 136] vs. 95 [89 – 99], p-value: 0.0005; 7 dpf: n = 8/8 DMSO/IOX1-treated animals; 153 [145 – 180] vs. 131 [129 – 137], p-value: 0.0033; 10 dpf: n = 8/8 DMSO/IOX1-treated animals; 192 [186 – 206] vs. 168 [150 – 173] oligodendrocytes, p-value: 0.0005; Wilcoxon rank sum test with Benjamini-Hochberg correction^57^). While the absolute number of of oligodendrocytes increased over time, the proportional decrease following IOX1 treatment was comparable across age (5 dpf: 22%; 7 dpf: 15%; 10 dpf: 13%).

Subcortical oligodendrocytes are unevenly distributed near balance nuclei. We therefore used a semi-automated analysis pipeline to quantify local differences in oligodendrocyte number in four balance-associated regions: the anterior MLF (Figure 1E), the lateral hindbrain (Figure 1F), the posterior MLF (Figure 1G), and the midbrain (Figure 1H). Confocal stacks from 7 dpf larvae expressing a fluorescent reporter in the cytoplasm of oligodendrocytes were first aligned to a reference stack^58^. Oligodendrocyte somata were manually marked and automatically counted (Methods) in regions of interest delineated in the reference stack. We observed reductions in oligodendrocyte numbers in the anterior MLF and lateral hindbrain (Figures 1E and 1F), but not the posterior MLF or the anterior midbrain (Figures 1G and 1H; oligodendrocytes DMSO vs. IOX1; n = 17/12 DMSO/IOX1-treated fish; anterior MLF: 8 [6 – 9] vs. 4 [3 – 5] adjusted p-value: 0.0077; lateral hindbrain: 26 [22 – 33] vs. 19 [16 – 22] adjusted p-value: 0.0084; posterior MLF: 20 [18 – 25] vs. 21 [20 – 24] adjusted p-value: 0.4110; Midbrain: 25 [18 – 28] vs. 18 [16 – 22] adjusted p-value: 0.1221; Wilcoxon rank sum test with Benjamini-Hochberg correction^57^). These regional differences suggest that balance circuit nodes might be differentially impacted by IOX1 treatment.

To evaluate whether the reduction in oligodendrocytes results in decreased myelination, we attempted to quantify the brightness of the fluorescent reporter signal under the mbp promoter in DMSO-treated control animals. We observed substantial variability among control fish in fluorescence in the anterior MLF (mean *±* SD: 5316 *±* 3965 au). Power analysis indicated that detecting a 15% reduction in myelin with *α* = 0.05 and *β* = 0.2 (power = 0.8) would require a sample size of 389 fish for each group. Given this impractical sample size, we did not continue with this analysis.

We performed two control experiments to investigate how IOX1 treatment reduces the number of oligodendrocytes, and to assay for potential off-target effects on balance neuron number. First, we measured the density of oligodendrocyte precursor cells after IOX1 treatment in the brains of 7 dpf larvae. Density was reduced (Figures S1F and S1G) indicating that IOX1 affects both proliferation and differentiation of oligodendrocyte precursor cells. Next, we counted neurons in three key balance populations: vestibular neurons in the tangential vestibular nucleus (TAN, Figure S1I), descending premotor neurons in the nucleus of the medial longitudinal fasciculus / interstitial nucleus of Cajal (nMLF/INC, Figure S1J), and extraocular motor neurons in the oculomotor and trochlear nuclei (nIII/nIV, Figures S1H and S1K). We observed no changes in the number of balance neurons after IOX1 treatment.

We conclude that early, continuous exposure to IOX1 disrupts oligodendrogenesis in select subcortical brain regions. The IOX1-induced decrease in oligodendrocytes persists across early development and does not affect the number of balance neurons in key midbrain/hindbrain circuits. Our model is therefore well-suited to explore the contributions of oligodendrocytes to circuit function as balance behaviors develop.

### IOX1 treatment impacts vertical navigation in an age-dependent manner

Gravity-guided vertical navigation is an important vestibular behavior^43^. Larval zebrafish navigate up/down by sequencing discreet movements that share a common trajectory (Figure 2A). To understand how disrupting oligodendrogenesis impacts vertical navigation, we measured kinematics and trajectories from freely-swimming zebrafish (Figure 2B) using our Scalable Apparatus to Measure Posture and Locomotion (SAMPL)^37^. All behavior was measured in the dark to isolate vestibular contributions to behavior. We compared the consistency of sequential trajectories (Figures 2C to 2E) in sets of IOX1-treated and control larvae at 4–6, 7–9, and 10–12 dpf (Figures 2F to 2H). In control fish, consistency was slightly higher from 4–6 than 7–9, but stabilized between 7–9 and 10–12 (DMSO 4 dpf: 0.85 [0.84 – 0.86] vs. 7 dpf 0.79 [0.78 – 0.79]; adjusted p-value < 0.0001; 7 dpf: 0.79 [0.78 – 0.79] vs 10 dpf: 0.78 [0.77 – 0.80] adjusted p-value: 0.4237).

**Figure 2:**
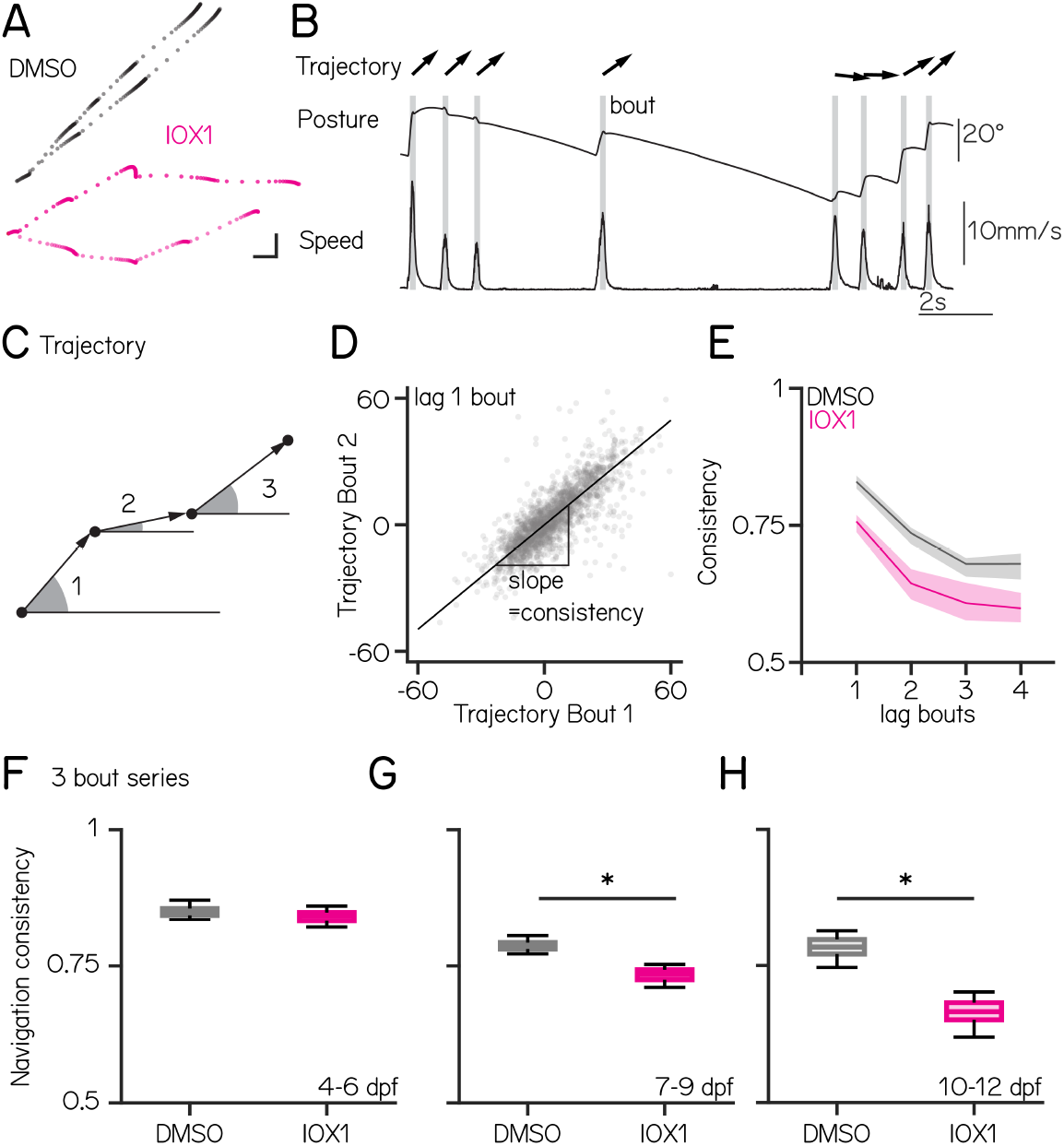
IOX1 treatment impacts vertical navigation in an age-dependent manner. **(A)** Fish position in time, plotted as dots, for two epochs containing multiple consecutive swim bouts from DMSO (gray) and IOX1-treated (pink) animals at 7 dpf. Scale bar 1mm **(B)** Graph of posture and swim speed for a single epoch. Individual swim bouts are highlighted in gray, arrows show trajectory. **(C)** Schematic of bout trajectories from three consecutive bouts illustrating the angle of the trajectory during peak speed. **(D)** Trajectory of bout 1 versus trajectory of bout 2 for all epochs from one control fish. Navigation consistency is defined as the slope of best fit line. **(E)** Navigation consistency as a function of swim bouts series for control (gray) and IOX1-treated fish (pink) at 7 dpf. **(F)** Average navigation consistency of three bout series for control and IOX1-treated fish at 4 – 6 dpf (DMSO vs. IOX1: 0.85 [0.84 – 0.86] vs. 0.84 [0.83 – 0.85] adjusted p-value: 0.466). **(G)** Average navigation consistency of three bout series for control and IOX1-treated fish at 7 – 9 dpf (DMSO vs. IOX1: 0.79 [0.78 – 0.79] vs. 0.73 [0.72 – 0.74] adjusted p-value: 0.002). **(H)** Average navigation consistency of three bout series for control and IOX1-treated fish at 10 – 12 dpf (DMSO vs. IOX1: 0.78 [0.77 – 0.80] vs. 0.67 [0.65 – 0.68] adjusted p-value: 0.0001). * indicates p-value < 0.05.

We observed an increasingly large impact of IOX1 treatment on navigation consistency with age. Despite having fewer oligodendrocytes at 5 dpf (Figure 1E), we did not detect any differences in navigation consistency at 4–6 dpf (Figure 2F; DMSO vs. IOX1: 0.85 [0.84 – 0.86] vs. 0.84 [0.83 – 0.85] adjusted p-value: 0.520). In contrast, IOX1-treated animals showed less consistent navigation from 7–9 (Figure 2G; DMSO vs. IOX1: 0.79 [0.78 – 0.79] vs. 0.73 [0.72 – 0.74] adjusted p-value: 0.002) and 10–12 dpf (Figure 2H, DMSO vs. IOX1: 0.78 [0.77 – 0.80] vs. 0.67 [0.65 – 0.68] adjusted p-value: 0.0016). We also analyzed a broad set of kinematic parameters for IOX1-treated fish. Little was different except that at 7 dpf, IOX1-treated fish had increased swim bout durations and reduced inter-bout intervals (Table 1). Finally, we observed no changes to navigation consistency in any of our control treatment groups: later, discontinuous, and lower doses of IOX1 (Figure S2 and Table 2).

**Table 1.** Behavioral results from 4 µM IOX1 treatment at 4, 7 and 10 dpf.

| Experiment | Metric | DMSO | IOX1 | p-value | p-value adj. |
| --- | --- | --- | --- | --- | --- |
| **4 dpf - 4 μM IOX1** | Navigation consistency | 0.85 [0.84–0.86] | 0.84 [0.83–0.85] | 0.279 | 0.520 |
|  | Inter-bout interval (IBI) [s] | 2.02 [1.63–3.10] | 1.77 [1.67–2.65] | 0.589 | 0.589 |
|  | IBI Pitch [°] | 12.39 [5.63–16.23] | 15.63 [11.23–15.99] | 0.485 | 0.554 |
|  | Speed [mm/s] | 9.71 [8.94–11.18] | 11.83 [10.85–12.33] | 0.041 | 0.164 |
|  | Duration [s] | 0.14 [0.14–0.16] | 0.16 [0.15–0.18] | 0.214 | 0.520 |
| **7 dpf - 4 μM IOX1** | Navigation consistency | 0.79 [0.78–0.79] | 0.73 [0.72–0.74] | 0.000 | 0.002 |
|  | IBI [s] | 2.97 [2.76–3.20] | 2.50 [2.12–2.68] | 0.005 | 0.012 |
|  | IBI Pitch [°] | 6.15 [4.16–7.02] | 7.71 [6.23–8.20] | 0.065 | 0.087 |
|  | Speed [mm/s] | 10.80 [9.24–11.68] | 11.61 [10.52–12.51] | 0.161 | 0.163 |
|  | Duration [s] | 0.16 [0.15–0.17] | 0.18 [0.17–0.18] | 0.007 | 0.014 |
| **10 dpf - 4 μM IOX1** | Navigation consistency | 0.78 [0.77–0.80] | 0.67 [0.65–0.68] | 0.000 | 0.000 |
|  | IBI [s] | 2.38 [2.11–3.25] | 1.80 [1.47–2.58] | 0.310 | 0.413 |
|  | IBI Pitch [°] | 3.69 [2.30–4.79] | 4.20 [2.43–5.96] | 0.690 | 0.690 |
|  | Speed [mm/s] | 9.56 [9.12–10.03] | 10.51 [8.86–11.06] | 0.548 | 0.626 |
|  | Duration [s] | 0.16 [0.16–0.17] | 0.15 [0.15–0.16] | 0.119 | 0.238 |

**Table 2.** Behavioral results for control experiments.

| Experiment | Metric | DMSO | IOX1 | p-value | p-value adj. |
| --- | --- | --- | --- | --- | --- |
| **7 dpf - 0.5 μM IOX1** | Navigation consistency | 0.77 [0.76–0.78] | 0.78 [0.77–0.79] | 0.344 | 0.688 |
|  | Inter-bout interval (IBI) [s] | 2.89 [2.77–3.10] | 2.71 [2.35–3.16] | 0.548 | 0.731 |
|  | IBI Pitch [°] | 5.02 [2.62–5.46] | 4.26 [3.87–5.37] | 1.000 | 1.000 |
|  | Speed [mm/s] | 10.35 [10.01–10.48] | 10.75 [10.28–11.53] | 0.222 | 0.688 |
|  | Duration [s] | 0.16 [0.15–0.17] | 0.16 [0.15–0.17] | 1.000 | 1.000 |
| **7 dpf - 2 μM IOX1** | Navigation consistency | 0.76 [0.75–0.77] | 0.80 [0.79–0.80] | 0.036 | 0.072 |
|  | IBI [s] | 3.01 [2.93–3.24] | 2.52 [2.39–2.63] | 0.026 | 0.069 |
|  | IBI Pitch [°] | 5.09 [4.57–5.76] | 5.42 [2.74–8.16] | 1.000 | 1.000 |
|  | Speed [mm/s] | 10.53 [9.64–11.46] | 10.75 [9.33–10.97] | 0.818 | 0.935 |
|  | Duration [s] | 0.16 [0.16–0.16] | 0.16 [0.16–0.17] | 0.431 | 0.575 |
| **7 dpf Early 4 μM IOX1** | Navigation consistency | 0.81 [0.80–0.82] | 0.78 [0.77–0.79] | 0.057 | 0.228 |
|  | IBI [s] | 2.17 [2.15–2.46] | 1.98 [1.83–2.35] | 0.200 | 0.320 |
|  | IBI Pitch [°] | 6.92 [4.38–7.69] | 6.59 [5.56–7.19] | 1.000 | 1.000 |
|  | Speed [mm/s] | 9.68 [8.47–11.35] | 9.51 [8.37–11.45] | 1.000 | 1.000 |
|  | Duration [s] | 0.16 [0.15–0.17] | 0.17 [0.16–0.18] | 0.314 | 0.419 |
| **7 dpf Late 4 μM IOX1** | Navigation consistency | 0.82 [0.81–0.82] | 0.83 [0.82–0.84] | 0.303 | 0.485 |
|  | IBI [s] | 2.64 [2.15–3.09] | 2.64 [2.43–2.94] | 1.000 | 1.000 |
|  | IBI Pitch [°] | 6.37 [4.64–7.22] | 4.96 [3.57–7.09] | 0.886 | 1.000 |
|  | Speed [mm/s] | 10.04 [9.00–11.01] | 9.68 [8.85–10.22] | 0.686 | 0.915 |
|  | Duration [s] | 0.15 [0.15–0.15] | 0.16 [0.16–0.17] | 0.029 | 0.077 |

We conclude that IOX1 treatment disrupts vertical navigation in an age-dependent manner.

### IOX1 treatment selectively impairs body tilt responses in nMLF/INC in older larvae

Impaired vertical navigation might reflect disrupted neuronal representation of body tilt relative to gravity. We focused on neurons in the TAN and the nMLF/INC as these two connected nuclei are key nodes in the circuit that transforms gravity-derived sensation into commands for vertical navigation^43^ (Figure 3A). To determine if IOX1 treatment impacts vestibular processing at 4 or 7 dpf (Figure 3B) in the nMLF/INC or the TAN, we used Tilt-In-Place Microscopy (TIPM; Figure 3C)^59^ to measure neuronal responses to body tilts. TIPM returns fish from a nose-up/nose-down orientation to the imaging plane faster (∼5 msec) than the time constant of the calcium indicator (GCaMP6s)^60^. The observed fluorescence reflects the decay of the neuron’s response to the body’s tilt.

**Figure 3:**
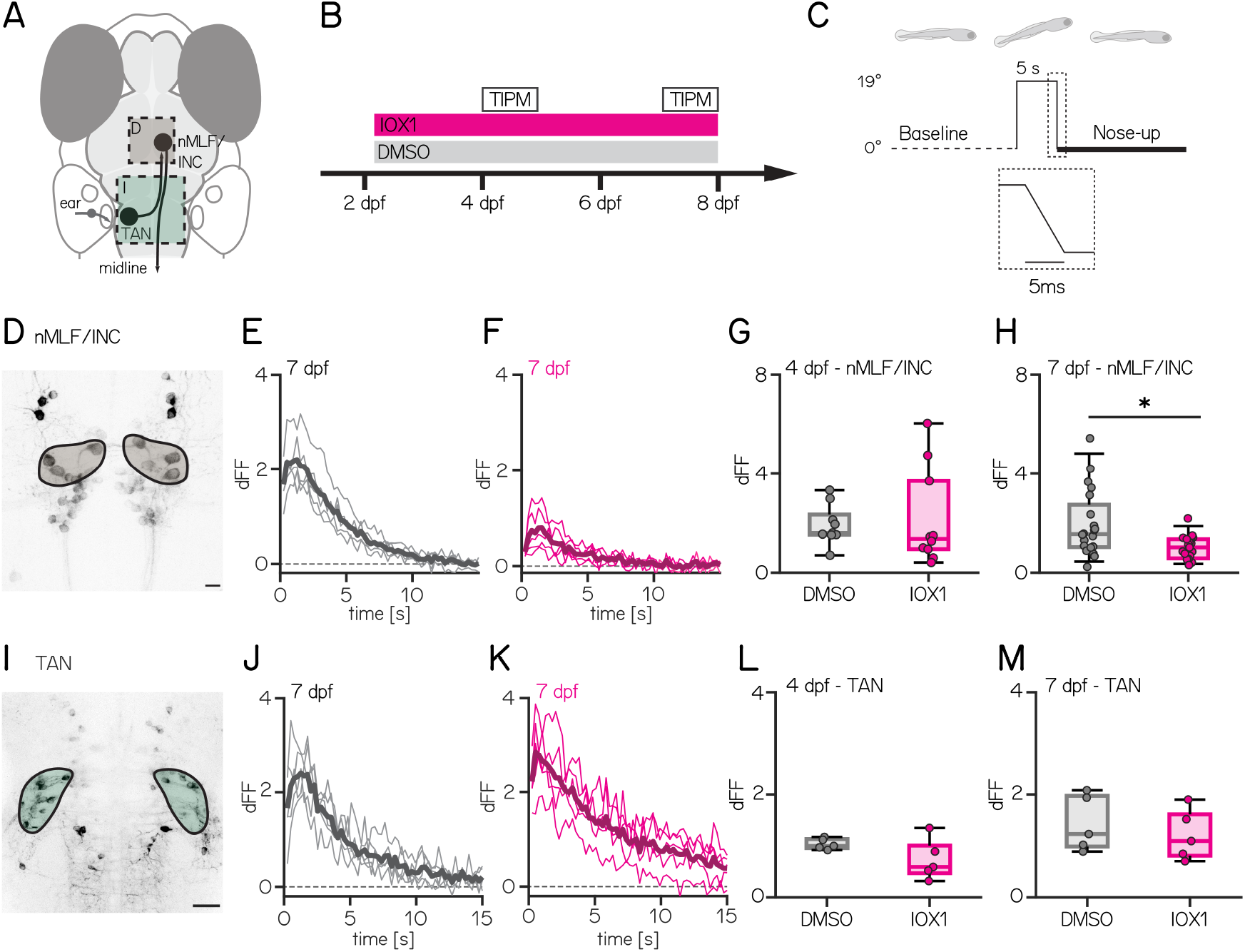
IOX1 treatment selectively impairs body tilt responses in nMLF/INC in older larvae. **(A)** Schematic of key nodes in the vertical navigation circuit, highlighting the tangential vestibular nucleus (TAN, green) which projects to the nucleus of the medial longitudinal fasciculus / interstitial nucleus of Cajal (nMLF/INC, brown). **(B)** Timeline of the experiment showing IOX1 and DMSO-only treatment timecourse beginning at 54 hours post-fertilization. Imaging timepoints are open rectangles at 4 and 7 days post-fertilization. **(C)** Timecourse of nose-up stimulus for imaging indicating pre-tilt baseline (dotted) and post-tilt imaging (thick line). Dotted rectangle corresponds to the 19° return to baseline. **(D)** Confocal image showing large spinal-projecting neurons in the nMLF/INC of the nMLF (brown) Scale bar: 10 µm **(E–F)** Normalized fluorescent changes after five nose-up tilts in DMSO (E, gray) and IOX1 (F, pink) from 7 dpf fish. Bold line is the median. **(G–H)** Average response from all neurons for each 4 (G) and 7 (H) dpf fish for DMSO (gray) and IOX1 (pink). (4 dpf: 1.60 [1.53 – 2.34] vs 1.35 [0.94 – 3.70], p-value: 0.7124; 7 dpf: 1.55 [1.03 – 2.74] vs 1.01 [0.58 – 1.35], p-value: 0.0355) **(I–M)** Same as (D–H) except for the tangential vestibular nucleus (TAN, green). (4 dpf: 1 [0.96 – 1.13] vs 0.59 [0.47 – 1.00], p-value: 0.3016; 7 dpf: 1.23 [0.99 – 1.97] vs 1.09 [0.81 – 1.61], p-value: 0.5476). **(I)** Scale bar: 20 µm. *indicates p-value < 0.05.

After IOX1 treatment, we observed an age-dependent decrease in responses to nose-up tilts in the nMLF/INC (Figure 3D). Neurons in 7 dpf IOX1-treated fish had a weaker response to 19° nose-up tilts (Figures 3E, 3F and 3H and Table 3; n = 20/20 DMSO/IOX1 treated animals: 1.55 [1.03 – 2.74] vs 1.01 [0.58 – 1.35], p-value: 0.0355). Consistent with the absence of behavioral effects from 4–6 dpf (Figure 2F) we saw no difference in responses in younger fish (Figure 3G; n = 10/10 DMSO/IOX1 treated animals; 1.60 [1.53 – 2.34] vs 1.35 [0.94 – 3.70], p-value: 0.7124). The nMLF/INC includes neurons that are morphologically distinct and identifiable across fish^52^. Of these, only the MeLr neurons had a significant decrease at 7 dpf, but we note that other neuron types (e.g. MeLc/MeLm) were less frequently labeled in our transgenic driver line (Table 4).

**Table 3.** nMLF/INC functional imaging (dFF)

| Age | Measurement | DMSO Median [25–75%] | IOX1 Median [25–75%] | p-value | p-value adj. |
| --- | --- | --- | --- | --- | --- |
| **4 dpf** | nMLF Up | 1.60 [1.53 – 2.34] | 1.35 [0.94 – 3.70] | 0.3154 | 0.7124 |
|  | nMLF IQR Up fish | 1.59 [0.80 – 2.25] | 0.91 [0.72 – 3.22] | 0.7802 | 0.8917 |
|  | nMLF IQR Up cell | 1.35 [0.81 – 2.31] | 0.92 [0.56 – 2.18] | 0.3092 | 0.7124 |
|  | nMLF Down | 0.77 [0.49 – 1.30] | 1.19 [0.75 – 3.44] | 0.1823 | 0.7124 |
|  | nMLF IQR Down fish | 1.19 [0.35 – 1.48] | 1.30 [0.75 – 2.62] | 0.3562 | 0.7124 |
|  | nMLF IQR Down cell | 1.06 [0.55 – 1.74] | 1.31 [0.57 – 1.89] | 0.4752 | 0.7603 |
| **7 dpf** | nMLF Up | 1.55 [1.03 – 2.74] | 1.01 [0.58 – 1.35] | 0.0093 | 0.0355 |
|  | nMLF IQR Up fish | 0.95 [0.60 – 1.49] | 0.56 [0.40 – 0.87] | 0.0543 | 0.0869 |
|  | nMLF IQR Up cell | 1.04 [0.67 – 1.73] | 0.49 [0.30 – 1.14] | 0.0002 | 0.0016 |
|  | nMLF Down | 0.81 [0.60 – 1.22] | 0.61 [0.36 – 0.86] | 0.0499 | 0.0869 |
|  | nMLF IQR Down fish | 0.59 [0.43 – 0.99] | 0.38 [0.30 – 0.58] | 0.0133 | 0.0355 |
|  | nMLF IQR Down cell | 0.54 [0.33 – 1.00] | 0.47 [0.27 – 0.76] | 0.1183 | 0.1577 |

**Table 4.** nMLF/INC cell type-specific analysis.

| **7 dpf** | **MeLr** | **MeLc** | **MeLm** | **MeM** |
| --- | --- | --- | --- | --- |
| n DMSO | 35 | 16 | 11 | 21 |
| dFF | 1.7 [1.1 – 3.3] | 1.3 [0.9 – 2.4] | 1.6 [0.6 – 4.1] | 1.1 [1.1 – 1.1] |
| n 4 μM IOX1 | 38 | 20 | 6 | 21 |
| dFF | 1.1 [0.5 – 1.7] | 0.9 [0.5 – 1.2] | 0.5 [0.4 – 0.6] | 0.3 [0.3 – 0.3] |
| p-value adj. | 0.0038 | 0.1286 | 0.2047 | 0.5 |

To better understand the decreased response of nMLF/INC neurons to body tilts, we repeated nose-up body tilts while imaging at an eccentric angle to characterize the full timecourse of the response^59^. There were no changes to response latency or decay in IOX1-treated fish at 7 dpf (Figure S3 and Table 5). We next evaluated spontaneous activity; we saw no change in frequency or amplitude at either 4 or 7 dpf in IOX1-treated fish (Figure S4). We conclude that the timecourse of tilt responses and basal activity levels in nMLF/INC neurons are unchanged after IOX1 treatment.

**Table 5.** nMLF/INC functional imaging at eccentric angles at 7 dpf.

| Age | Measurement | DMSO Median [25–75%] | IOX1 Median [25–75%] | p-value | p-value adj. |
| --- | --- | --- | --- | --- | --- |
| **7 dpf** | peak time [s] | 2.3 [1.5 – 3.6] | 2.0 [1.3 – 2.6] | 0.4394 | 0.6591 |
|  | decay slope [dFF/s] | -0.003 [-0.025 – 0.004] | -0.008 [0.022 – -0.002] | 0.7104 | 0.7104 |

To evaluate whether prolonged exposure to IOX1 affected neuronal responses, we evaluated nMLF/INC responses in larvae that began IOX1 treatment at 72 hpf, instead of 54. Despite receiving over 4 full days of IOX1 treatment, we did not observe any difference in nMLF/INC responses between control and later-treated animals (Figure S5, DMSO vs. IOX1 nose-up responses: 1.82 [0.98 – 2.67] vs. 2.99 [2.69 – 3.98], p-value: 0.0539; nose-down responses: 0.71 [0.64 – 0.89] vs. 1.15 [0.87 – 1.91], p-value; 0.0539). We did not see any changes in the number of oligodendrocytes under the same delayed IOX1 treatment protocol (Figure S1C). We therefore conclude that prolonged exposure to IOX1, if delayed relative to oligodendrogenesis, is insufficient to disrupt vestibular processing.

In contrast to the nMLF/INC, after IOX1 treatment, neurons in the TAN did not respond differentially to body tilts (Figures 3J and 3K). Responses to nose-up tilts in treated animals and controls were indistinguishable at both 4 and 7 dpf (Figures 3L and 3M and Table 6; DMSO vs IOX1 4 dpf: 1 [0.96 – 1.13] vs 0.59 [0.47 – 1.00], p-value: 0.3016; 7 dpf: 1.23 [0.99 – 1.97] vs 1.09 [0.81 – 1.61], p-value: 0.5476). Finally, we saw no differences between control and IOX1-treated fish at any age to nose-down tilts in either the nMLF/INC or TAN (Figure S6). While we cannot exclude that we missed the “true” nose-down neurons due to restricted expression in our driver line^61^, many of the neurons we measured did respond to nose-down stimuli, just with markedly smaller changes in fluorescence (dFF < 2).

**Table 6.** TAN functional imaging at 4 & 7 dpf.

| Age | Measurement | DMSO Median [25–75%] | IOX1 Median [25–75%] | p-value | p-value adj. |
| --- | --- | --- | --- | --- | --- |
| **4 dpf** | TAN 4 dpf Up | 1 [0.96 – 1.13] | 0.59 [0.47 – 1.00] | 0.1508 | 0.3016 |
|  | TAN 4 dpf Down | 1.15 [0.96 – 1.36] | 0.55 [0.47 – 1.40] | 0.4206 | 0.4206 |
| **7 dpf** | TAN 7dpf Up | 1.23 [0.99 – 1.97] | 1.09 [0.81 – 1.61] | 0.4206 | 0.5476 |
|  | TAN 7 dpf Down | 1.27 [0.98 – 1.44] | 1.10 [0.74 – 1.30] | 0.5476 | 0.5476 |

Taken together our data support the hypothesis that disrupted oligodendrogenesis following IOX1 treatment decreases the sensitivity of select nMLF/INC neurons to nose-up tilts. Furthermore, central vestibular representations of body tilt are preserved.

### IOX1 treatment does not change vestibulo-ocular reflex behavior or extraocular motor neuron responses

Information about body tilt magnitude encoded by TAN neurons is relayed (Figure S7A) to extraocular motor neurons in cranial nuclei nIII/nIV (Figure S7B) by means of a branch off the TAN axon that projects to the nMLF/INC^49^. This projection is responsible for the vestibulo-ocular reflex (Figure S7C), that stabilizes gaze by counter-rotating the eyes after nose-up/nose-down body tilts^45^. If central vestibular representations of body tilt are preserved following IOX1 treatment then gaze stabilizing behavior and extraocular motor neuron responses should be unaffected. We evaluated the nose-up/eyes-down vestibulo-ocular reflex and performed TIPM in nIII/nIV. IOX1 treatment did not change the strength of the vestibulo-ocular reflex at either 4 or 7 dpf (Figures S7D to S7F and Table 7). We conclude that IOX1 treatment does not significantly impact the TAN, its extraocular motor neuron targets, and the gaze-stabilizing behavior they subserve.

**Table 7.** nIII and nIV motor neuron functional imaging and vestibulo-ocular reflex performance at 4 and 7 dpf.

| Age | Measurement | DMSO Median [25–75%] | IOX1 Median [25–75%] | p-value | p-value adj. |
| --- | --- | --- | --- | --- | --- |
| **4 dpf** | nIII 4 dpf Up [dFF] | 0.84 [0.65 – 0.97] | 0.72 [0.53 – 1.17] | 0.6607 | 0.8809 |
|  | nIV 4 dpf Up [dFF] | 0.90 [0.65 – 1.46] | 0.92 [0.48 – 1.47] | 1.0000 | 1.0000 |
|  | VOR gain 4 dpf | 0.81 [0.63 – 0.84] | 0.58 [0.46 – 0.95] | 0.7209 | 0.7209 |
| **7 dpf** | nIII 7 dpf Up [dFF] | 0.59 [0.50 – 0.68] | 0.82 [0.44 – 1.2] | 0.4208 | 0.4809 |
|  | nIV 7 dpf Up [dFF] | 1.20 [0.87 – 1.54] | 0.66 [0.50 – 2.36] | 0.6764 | 0.6764 |
|  | VOR gain 7 dpf | 0.59 [0.45 – 0.92] | 0.72 [0.62 – 1.35] | 0.3660 | 0.3660 |

Taken together, our results argue that vestibular processing — both peripheral and central — and gaze-stabilization behavior are not affected by IOX1 treatment. These controls therefore highlight the susceptibility of the nMLF/INC node in the vertical navigation circuit.

### Photoablation of oligodendrocytes near the anterior MLF disrupts vertical navigation

Taken together, our results support a model where interfering with oligodendrogenesis in the anterior MLF (Figure 4A) disrupts vertical navigation in older larvae. We tested this hypothesis by acutely photoablating all of the oligodendrocytes whose cell bodies sat between the left and right branches of the MLF (Figures 4B and 4D and Table 10, 4 dpf: 75 [64 – 95]% vs 6 dpf 83 [61 – 90]%; p-value: 0.6307). One day after photoablation we observed significantly reduced myelin along the MLF (Figures 4B and 4C, Control vs. lesion: 1.07 [1 – 1.09] vs. 0.56 [0.51 – 0.66], p-value: 0.0357). As with IOX1 treatment, we saw no disruption to vertical navigation on days 5–6 after photoablation of cells at 4 dpf (Figure 4E and Tables 8 and 10, navigation consistency control vs. lesion: 0.89 [0.88 – 0.90] vs. 0.85 [0.83 – 0.86], p-value: 0.272). Importantly, photoablation at 6 dpf led to a significant decrease in navigation consistency at 7–9 dpf (Figure 4F, 0.83 [0.82 – 0.84] vs. 0.75 [0.74 – 0.76], p-value: 0.008). We infer that demyelination of the anterior MLF after photoablation of nearby oligodendrocytes selectively disrupts vertical navigation in older larvae.

**Table 8.** Behavioral analysis at 5 and 7 dpf after oligodendrocyte lesions in the anterior MLF.

| Condition | Metric | Ctrl Median [25-75%] | Demyelination Median [25-75%] | p-value | p-value adj. |
| --- | --- | --- | --- | --- | --- |
| 4 dpf Demyelination | Navigation consistency | 0.89 [0.88-0.90] | 0.85 [0.83-0.86] | 0.068 | 0.272 |
|  | IBI [s] | 2.21 [2.16-2.36] | 1.93 [1.78-2.71] | 0.700 | 0.914 |
|  | IBI Pitch [°] | 7.59 [5.64-16.27] | 5.21 [3.53-6.85] | 0.400 | 0.640 |
|  | Speed [mm/s] | 10.76 [10.40-11.18] | 10.30 [9.84-11.28] | 1.000 | 1.000 |
|  | Duration [s] | 0.14 [0.14-0.15] | 0.15 [0.15-0.15] | 0.800 | 0.914 |
| 7 dpf Demyelination | Navigation consistency | 0.83 [0.82-0.84] | 0.75 [0.74-0.76] | 0.001 | 0.008 |
|  | IBI [s] | 1.43 [1.29-1.55] | 1.94 [1.66-2.94] | 0.200 | 0.320 |
|  | IBI Pitch [°] | 6.89 [5.31-10.13] | 3.52 [2.60-4.36] | 0.100 | 0.200 |
|  | Speed [mm/s] | 14.62 [13.27-15.73] | 12.93 [12.72-15.47] | 1.000 | 1.000 |
|  | Duration [s] | 0.16 [0.14-0.16] | 0.15 [0.14-0.15] | 1.000 | 1.000 |

**Table 9.** Numbers of cells imaged with TIPM for different experiments, nuclei, and ages.

| Experiment | DMSO Median [25–75%] | IOX1 Median [25–75%] | n cells DMSO | n cells IOX1 | n fish DMSO | n fish IOX1 |
| --- | --- | --- | --- | --- | --- | --- |
| 4 dpf nMLF/INC | 5 [4 – 6] | 5 [3 – 5] | 52 | 45 | 10 | 10 |
| 7 dpf nMLF/INC | 4 [3 – 5] | 4 [3 – 6] | 83 | 85 | 20 | 20 |
| 7 dpf nMLF/INC late | 6 [5 – 6] | 6 [5 – 6] | 56 | 56 | 10 | 10 |
| 7 dpf nMLF/INC eccentric | 5 [3 – 6] | 5 [5– 5] | 32 | 30 | 7 | 7 |
| 4 dpf TAN | 19 [17 – 22] | 19 [17 – 21] | 96 | 96 | 5 | 5 |
| 7 dpf TAN | 23 [18 – 25] | 27 [20 – 29] | 107 | 126 | 5 | 5 |
| 4 dpf nIII | 8 [6 – 13] | 4 [3 – 6] | 96 | 45 | 10 | 10 |
| 7 dpf nIII | 6 [4 – 8] | 5 [2 – 7] | 120 | 93 | 20 | 20 |
| 4 dpf nIV | 5 [2 – 8] | 5 [3 – 5] | 49 | 42 | 10 | 10 |
| 7 dpf nIV | 5 [2 – 6] | 1 [0 – 5] | 90 | 48 | 20 | 20 |

**Table 10.** Quantification of ablated oligodendrocytes.

| Experiment | Quantification | Demyelination |
| --- | --- | --- |
| 7 dpf for myelin quantification | lesioned cells | 6 [6 – 7] |
|  | % Oligodendrocyte loss in anterior MLF | 83 [67 – 86] |
| 4 dpf pre behavior | lesioned cells | 4 [3 – 4] |
|  | % Oligodendrocyte loss in anterior MLF | 75 [64 – 96] |
| 7 dpf pre behavior | lesioned cells | 5 [5 – 7] |
|  | % Oligodendrocyte loss in anterior MLF | 83 [61 – 90] |

**Figure 4:**
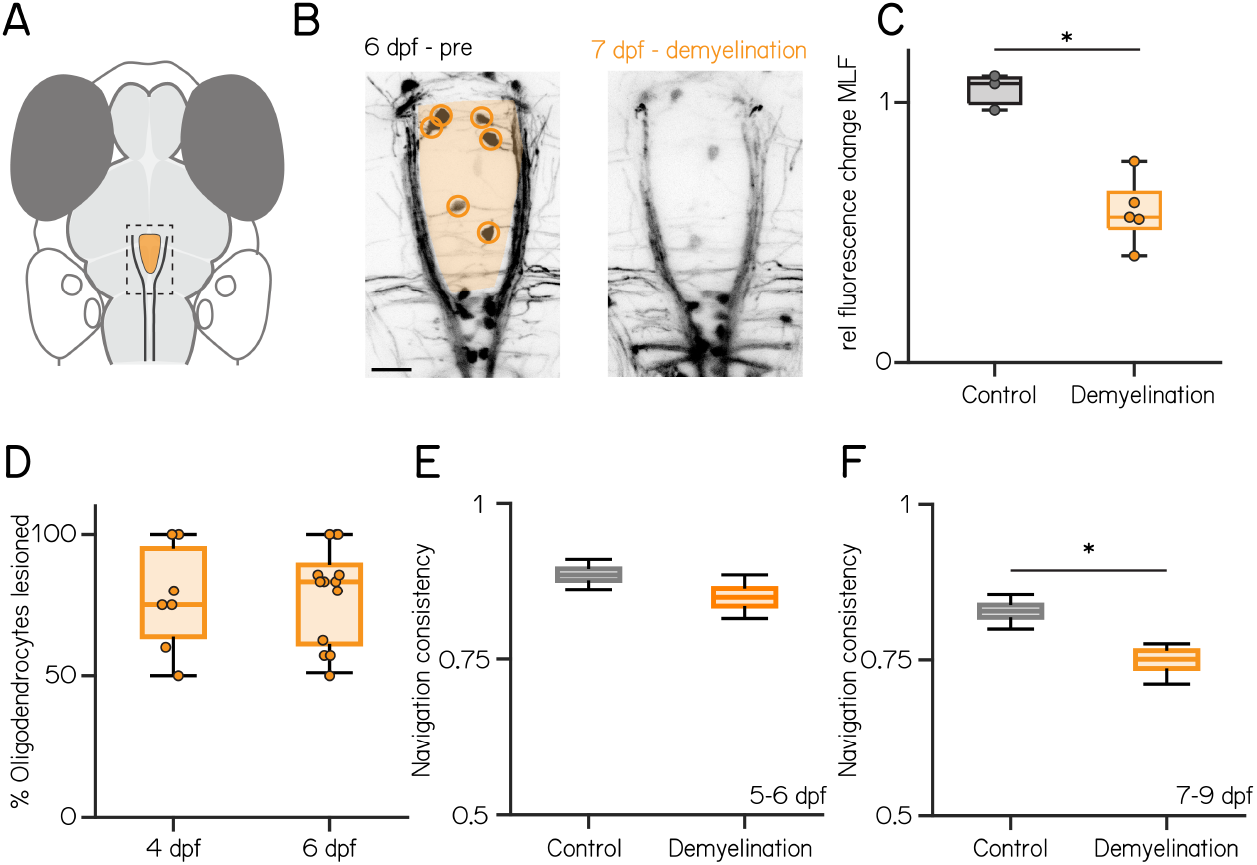
Photoablation of oligodendrocytes near the anterior MLF disrupts vertical navigation in older fish. **(A–B)** Schematic and confocal images of the anterior MLF before (6 dpf, left) and 1 day after oligodendrocyte lesion. Scale bar: 10 µm. **(C)** Relative fluorescence change in the MLF in control (gray) and lesioned fish (Control vs. lesion: 1.07 [1 – 1.09] vs. 0.56 [0.51 – 0.66], p-value: 0.0357). **(D)** Fraction of lesioned oligodendrocytes per animal at 4 or 6 dpf (4 dpf: 75 [64 – 95]% vs 6 dpf 83 [61 – 90]%; p-value: 0.6307), **(E–F)** Navigation consistency in control (gray) and lesioned (yellow) fish from 5–6 (E) and 7–9 (F) dpf (5–6 dpf: 0.89 [0.88 – 0.90] vs. 0.85 [0.83 – 0.86], p-value: 0.272; 7–9 dpf: 0.83 [0.82 – 0.84] vs. 0.75 [0.74 – 0.76], p-value: 0.008). indicates p-value < 0.05.

## DISCUSSION

We demonstrate behaviorally-specific, localized, and age-dependent consequences for balance following disrupted oligodendrogenesis. IOX1 treatment consistently decreased the number of oligodendrocytes in balance-relevant regions of the midbrain and hindbrain across early development. Despite the consistent decrease, IOX1 treatment disrupted vertical navigation in an age-dependent manner: young larvae were not affected, and deficits became more pronounced as larvae matured. Similarly, midbrain vestibulo-recipient neurons in the nMLF/INC were less responsive to postural stimuli, but only in older larvae. However, both the responses of upstream hindbrain vestibular neurons and a second gravity-dependent behavior, vertical gaze-stabilization, were unaffected. Acute photoablation of oligodendrocytes in the anterior MLF led to local demyelination and confirmed our behavioral findings, disrupting vertical navigation in older, but not younger, larvae. These experiments show the emergent importance of oligodendrocytes for proper postural encoding by nMLF/INC neurons. Taken together, our work reveals that the earliest functional contributions of oligodendrocytes can be remarkably restricted, impacting particular neurons at particular times as balance behaviors mature.

Oligodendrogenesis^14–16,62^ supports motor learning in mature animals. Do new oligodendrocytes similarly shape maturation of vertical navigation? In control larvae, vertical navigation consistency changes between 4–6 (when fish begin to swim) and 7–9 dpf; we see no change between 7–9 dpf and 10–12 dpf. IOX1-treated fish had consistently fewer oligodendrocytes from 5–10 dpf. As oligodendrogenesis is disrupted before and throughout maturation of vertical navigation, studies of adult motor learning anticipate a similarly persistent behavioral effect. Instead, we saw no change to vertical navigation from 4–6 dpf, and an increasingly pronounced impact from 7–9 to 10–12 dpf. Further, acute loss of oligodendrocytes was sufficient to disrupt behavior at 7–9 dpf. The early dispensability and acute inducibility of our effects suggest that the contribution of oligodendrocytes increases with time, more closely resembling classical models of a progressive increase in circuit capacity that follows myelination^63,64^.

Oligodendrogenesis and myelination also shape cortical critical periods: times during development of heightened sensitivity and neural plasticity^26,30–32^. Unexpectedly, given that demyelination of the MLF disrupts the adult vestibuloocular reflex^65^, IOX1 treatment did not disrupt the vestibulo-ocular reflex, the vestibular hindbrain neurons responsible^45,49^, or the relevant extraocular motor neurons. Although the number of hindbrain oligodendrocytes near TAN neurons decreased, we cannot rule out the possibility that IOX1-mediated changes are too small to affect the vestibuloocular reflex. Alternatively, the vestibulo-ocular reflex is different from critical period-associated behaviors in that it must remain plastic throughout life to adjust to growing and aging sensory and motor machinery^8,66^. Further, unlike the cognitive/behavioral disruption that follows social isolation or the amblyopia that follows monocular deprivation, sensory feedback (visual and vestibular) are both dispensable for the maturation of the vestibulo-ocular reflex^46^. In the context of this study, the robustness of the vestibulo-ocular reflex circuit to IOX1 treatment provides an important control for non-specific effects of pharmacological manipulation. More broadly, our finding that the early vestibulo-ocular reflex circuit is robust to disrupted oligodendrogenesis — even when the mature behavior relies on myelinated axons, and vertical navigation is impacted — expands our models of the role of oligodendrocytes in sensorimotor circuit development.

The oligodendrocytes whose cell bodies are proximal to the anterior MLF myelinate two sets of axons. The first population are “input” axons that originate from cells in the TAN and project to the nMLF/INC and nIII/nIV^45,49,67^. TAN neurons in IOX1-treated fish had normal responses to body tilts, and IOX1-treated fish had a normal vestibulo-ocular reflex. The second are “output” axons that descend to the spinal cord from cells in the nMLF/INC^43,68–70^. Only nMLF/INC neurons showed decreased responses to body tilts after IOX1 treatment or photoablation of anterior MLF oligodendrocytes, consistent with impaired vertical navigation. Our experiments therefore suggest that anterior MLF oligodendrocytes that myelinate the proximal region of nMLF/INC axons play a privileged role in maturation of vertical navigation.

Classically, myelination is associated with increased conduction velocity^71^, and demyelination of spinal axons is associated with balance deficits^72^. The small size of the larval zebrafish and seconds-long timescale of navigation behavior suggest that if the consequences we see after disrupting oligodendrocytes follow from changes to myelin, the deficits after photoablation are unlikely to follow from decreased conduction velocity. As their TAN-derived input, response latency, and response profile are all unchanged, we hypothesize that instead, the changes we see reflect an inability of nMLF/INC neurons to generate action potentials at high frequencies. Hypomyelination of a descending neuron in the larval zebrafish can interfere with action potential generation^73^, and excitability changes after hypo- or demyelination^74–78^. Future electrophysiological recordings of the responses of nMLF/INC and TAN neurons to current injection after oligodendrocyte photoablation and/or IOX1 treatment could address this directly.

In conclusion, we propose that the combination of different treatment schemes, concentrations, and targeted demyelination experiments convincingly demonstrates that loss of anterior MLF myelin disrupts nMLF/INC neuron function. The preservation of vestibular neuron function and vestibulo-ocular reflex behavior support a model where the inputs to nMLF/INC are preserved, but processing those inputs becomes progressively compromised with age. Consequentially, larvae with disrupted oligodendrogenesis show increasingly severe impacts on navigation, but not to gaze stabilization. Oligodendrocytes are essential for proper development of neural circuits and behaviors; failure results in severe neurological symptoms^79,80^. By dissociating where and when developmentally-disrupted oligodendrogenesis impacts neuronal function and behavior, our work takes a major step towards a mechanistic understanding of the contribution of oligodendrocytes to the maturation of sensorimotor behaviors.

## MATERIALS AND METHODS

### Fish care

The Institutional Animal Care and Use Committee of New York University Grossman School of Medicine approved all procedures involving zebrafish (*Danio rerio*) larvae. Fertilized eggs were collected and maintained at 28.5° C on a standard 14/10 hour light/dark cycle at densities of 20–50 larvae per 10 cm diameter petri dish. Dishes were filled with 25–40 mL E3 medium with 0.5 ppm methylene blue for the first 24–48hrs and then changed to E3 medium alone. After 5 dpf, larvae were maintained at densities under 20 larvae/dish and fed cultured rotifers (Reed Mariculture) daily.

### Transgenic zebrafish lines

We used the following published lines of transgenic zebrafish: *Tg(mbpa:NLS-EGFP)*^*zf3078Tg*^^81^; *Tg(mbpa:KillerRed)*^25^; *Tg(nefma:hsp70l-LOXP-RFP-LOXP-GAL4)*^*stl601Tg*^^82^; *Tg(6*.*7Tru*.*Hcrtr2:GAL4-VP16)*^*a150Tg*^^83^; *Tg(14xUAS:GCaMP6s)*^*mpn101Tg*^^68^; and *Tg(isl1a:GFP)*^*rw0Tg*^^84^.

### IOX1 treatment

IOX1 (Sigma Aldrich SML0067) was diluted in DMSO (Sigma Aldrich D8418) to a 10 mM stock solution. The stock solution was stored at −20° C. The treatment solution was diluted fresh on the day of use a final DMSO concentration of 0.2% in E3 embryo medium. For control treatments 0.2% DMSO in E3 was used. Drug or DMSO solutions were replaced on day 5 and day 7. IOX1 treatment started at 54 hours post fertilization (hpf) for normal treatment and at 72 hpf for late treatment start. Fish were kept in drug solution until the end of the experiment except for the early treatment group where fish were put in E3 medium at 72 hpf.

### Oligodendrocyte photoablation

Oligodendrocytes were visualized in *Tg(mbpa:KillerRed)* fish using a microscope (Thorlabs Bergamo) equipped with two pulsed infrared lasers: one for imaging with a high repetition rate (either a Spectra-Physics Mai Tai HP or a Coherent Axon) and one with high power and a low-repetition rate (Spectra-Physics Spirit 8W) for cell ablation. Fish were mounted in 2% low-melting temperature agarose, and imaged to define single-pixel regions of interest within oligodendrocytes. Photoablation was performed by exposing the region of interest to 1–4 laser pulses (1040 nm, 35–50 nJ at the sample, 400 fs). Ablation was confirmed by the absence of or disaggregated fluorescence in subsequent images of the targeted oligodendrocyte. Demyelination was quantified on confocal stacks (Zeiss LSM 800, 20x/1.0 NA objective) taken before and 1 day after photoablation. Regions of interest were drawn along the the demyelinated tracts (excluding any somata) and average intensity computed using Fiji^85^.

### Oligodendrocyte quantification

To count oligodendrocytes in specific regions, confocal stacks of *Tg(mbpa:KillerRed)* fish were first aligned to a reference stack using Fiji^85^ BigWarp^58^. We used 20 – 30 landmarks using the Rotation or Similarity warping method depending on the quality of the final overlap with the reference stack (qualitative assessment by experimenter). After alignment, the oligodendrocyte locations were marked using the CellCounter tool in Fiji. Regions of interest for each brain region were defined on a projection of the reference stack. A custom MATLAB (Mathworks, Natick MA, USA) script was used to import oligodendrocyte coordinates and brain ROIs and to report the number of oligodendrocytes in each region.

### Postural behavior

Behavior was measured using the Scalable Apparatus for Measuring Posture and Locomotion (SAMPL) that consists of a chamber where larvae could swim freely, an infrared illuminator, a camera, and software to process video in real time. A comprehensive description of the apparatus, processing algorithms, and baseline data for different genetic backgrounds can be found in^37^. Briefly, larvae were transferred to chambers at densities of 3–4 fish per chamber for 7 dpf experiments or 1–4 fish per chamber for 10 dpf experiments. The chambers contained 25–30 ml of 0.2% DMSO in E3 for controls, and 0.5, 2, or 4 µM IOX1 in 0.2% DMSO. Fish from each clutch (siblings) were randomly assigned to control and treatment groups. Data from fish from the same clutch and condition were pooled to form one experimental repeat (minimum 2–3 apparatus / condition). Only fish with inflated swim bladders were used. For the 7 and 10 dpf behavior experiments, behavior recordings were paused after 24 h for 30–60 minutes during which 1–2 ml of rotifer culture was added to each chamber. Larvae were removed from the apparatus after 48 h.

The number of analyzed bouts, animals per experiment, and experimental repeats are documented in Table 11. Here, navigation consistency is defined as the average of the slopes of the best fit line (bisquare regression model) of the trajectory of the first bout of a series against the trajectory of the 2nd and 3rd bout of a series. Only bout series with an average inter-bout interval of at least 2 seconds Figure S2A were used to compute navigation consistency. Behavior data was analyzed using custom-written software in MATLAB to extract individual swim bouts from the raw data (x/z position and pitch angle as a function of time). Only bouts during the circadian day were analyzed.

**Table 11.**
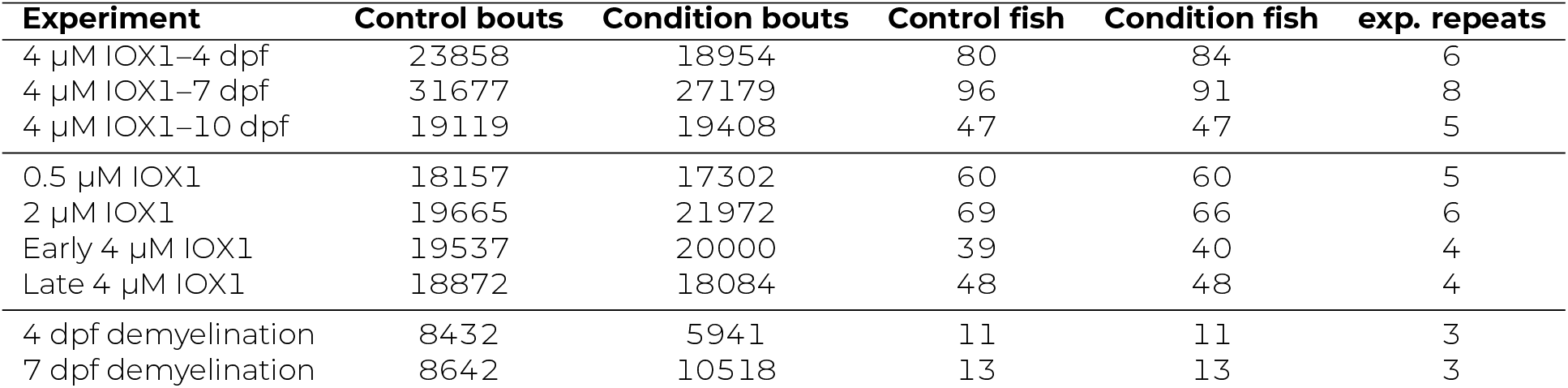
Analyzed bout numbers, fish numbers, and experimental repeats for behavioral experiments.

### Functional imaging

Fish were screened prior to drug treatment for GCaMP6s expression and then randomly split into control and treatment groups. Fish were kept in drug or control solution until they were mounted for imaging. All calcium imaging experiments were performed using Tilt In Place Microscopy (TIPM), described comprehensively in^59^. Briefly, 4 or 7 dpf fish were mounted in the center of the uncoated side of a mirror galvanometer (GVS0111, Thorlabs) in 2% low-melting-point agarose. The agarose around the tail was removed to facilitate exposure to the drug or DMSO solution. E3 was placed over the agarose, and the galvanometer mirror was placed under the microscope with a 20x/1.0 NA objective. A microscope (Thorlabs Bergamo) was used to measure fluorescence elicited by multiphoton excitation using a pulsed infrared laser (Mai Tai HP and Axon, both at 920 nm). Fast volumetric scanning was achieved using a piezo actuator (PFM450E, Thorlabs) to move the objective. Each frame of the volume (192 x 96 pixels) was collected with a 0.6 µs pixel dwell time (19.67 frames/s) resulting in a sampling rate of 3.93 volumes/s. Each fish was presented with 5 trials consisting of: 1) a 15 sec baseline period 2) a 5 sec −19° stimulus 3) a 15 sec recording period 4) a 5 sec 19° stimulus and 5) a 15 sec recording period. Cell identity in the nMLF/INC and nIII/nIV was determined based on morphology and location in an anatomy stack. The number of cells imaged for the different nuclei and ages are reported in Table 9. Regions of interest drawn in Fiji were loaded into MATLAB to extract the intensity of fluorescence after motion correction was performed^86^. Tilt responses are defined as the maximum normalized change in fluorescence (dFF) during the the first second of the recording period. For eccentric imaging 9 trials following 19° nose-up were recorded for each fish. To estimate the baseline fluorescence for each cell, 0.2mg/ml tricaine was added to anesthetize the fish. 3 more baseline trials with 19° nose-up stimuli were recorded. The time of peak dFF and the slope of the GCaMP6s signal during the last 2/3 of the stimulus were analyzed.

### Eye movement recordings

Torsional eye rotations were measured in response to step tilts delivered as described in^46^. Briefly, fish were mounted in 2% low-melting temperature agarose. Agarose around the left eye was removed, and the fish were placed in a glass cuvette on a rotating platform that also contained an IR illuminator and a camera focused on the eye. The platform was then rotated for 50 cycles of 15° steps (peak velocity 35°/sec, peak acceleration of 150°/sec^2^). Eye rotation was extracted from video frames acquired at 200 Hz using a pattern matching algorithm. The eye’s rotation following each step was evaluated manually; steps with rapid deviations in eye position indicative of horizontal saccades or gross failure of the tracking algorithm were excluded from analysis. The gain was defined as the ratio of peak eye velocity to peak table velocity within the first second after step onset. For each fish, the median gain across trials was used as the representative value.

### Statistics

All statistical testing was performed in MATLAB R2024b. All kinematic parameters were analyzed by experimental repeat except for fin-body coordination and navigation consistency, which were pooled across repeats to allow sufficient sampling. To estimate the spread of the data for navigation consistency, we resampled (100 times, with replacement) from the data from each condition and computed the expected value for control and perturbed datasets. This bootstrapped dataset was used to explicitly compute a p-value for fitted variables. The minimum number of bouts for navigation consistency was estimated using increasing sample sizes and fitting the variables until the coefficient of variation fell below 2%.

For all other parameters we performed two-sided Wilcoxon rank sum tests. To correct for multiple comparisons, the p-values were adjusted using the Benjamini-Hochberg correction^57^ with a false discovery rate of 0.05. The calculated and adjusted p-values are listed in the supplemental tables (Tables 1 to 3 and 5 to 8). All data are presented as median and the bounds of the inter-quartile range [25% – 75%]. This led to a minimum number of bout series of 1000.

For linear fits a robust regression model (bisquare) was used. Because the p value was computed explicitly and not estimated with respect to a theoretical probability distribution, we did not report an N for fitted variables (Navigation consistency).

## Data & Code

All data, raw and analyzed, as well as code necessary to generate the figures is available at 10.17605/OSF.IO/7UH2Y

## ACKNOWLEDGMENTS

The authors would like to thank Kristen Severi for help with identifying nMLF/INC neurons. Research was supported by the National Institute on Deafness and Communication Disorders of the National Institutes of Health under award number R01DC017489 to DS and K99DC021729 to FA, and a Hearing Health Foundation grant to FA. The authors would like to thank James Salzer, Bradley Zuchero, Yunlu Zhu, Celine Bellegarda, Louise Schenberg, Samantha David, Katherine Nagel, and members of the Schoppik and Nagel labs for their valuable feedback and discussions.

## AUTHOR CONTRIBUTIONS

Conceptualization: FA and DS, Methodology: FA, Investigation: FA and YZ, Visualization: FA, Writing: DS and FA, Editing: DS, Funding Acquisition: FA and DS, Supervision: DS and FA.

---

**Figure S1:**
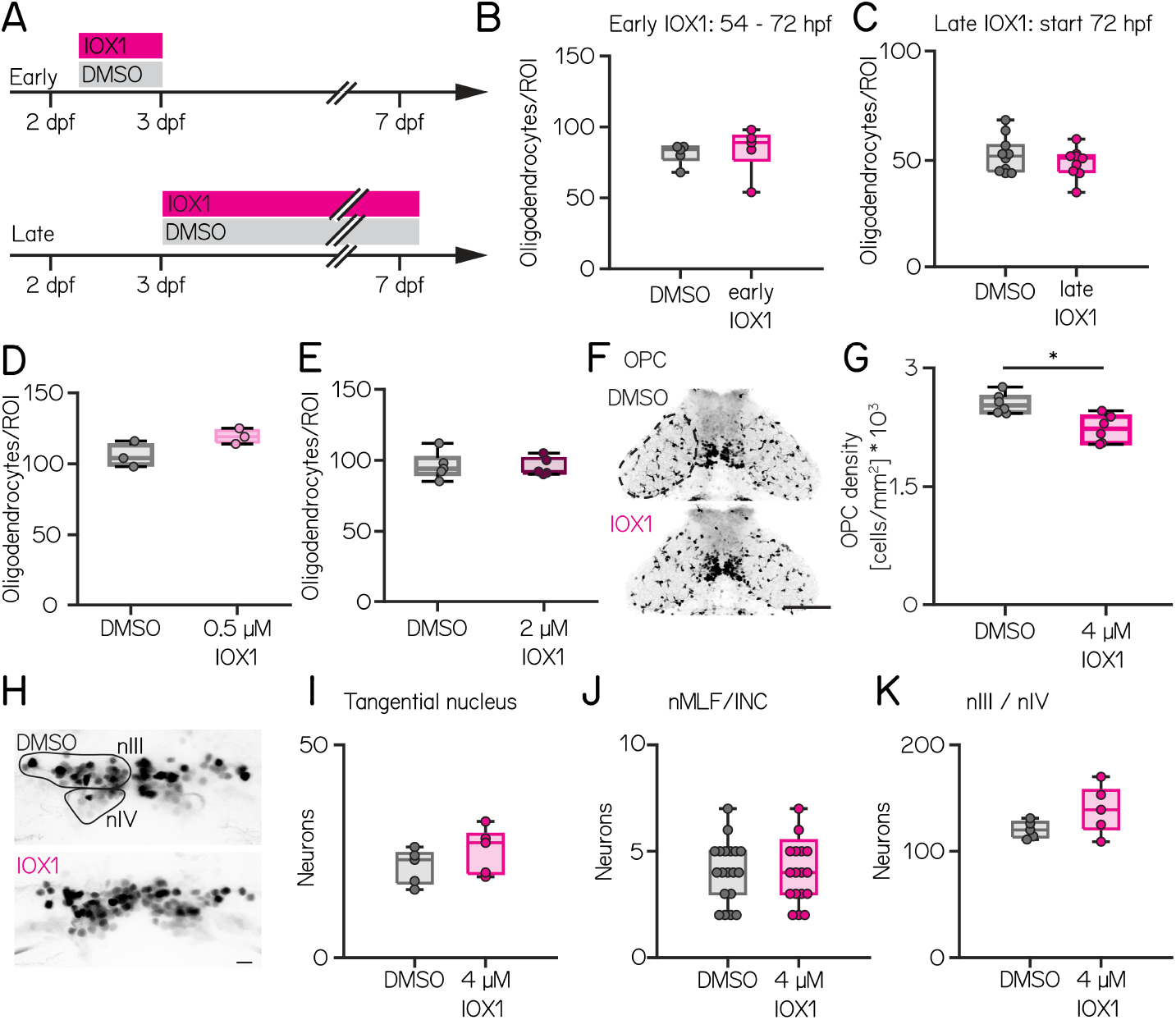
Different timing or concentrations of IOX1 do not affect oligodendrocyte numbers. **(A)** Schematic of early (top) and late (bottom) 4 µM IOX1 treatment. White square indicating the analysis timepoint at 7 dpf. **(B)** Oligodendrocyte counts at 7 dpf in control (gray) and 4 µM early IOX1-treated larvae (0.81 [0.80–0.82] vs 0.78 [0.77–0.79]; p-value = 0.228). **(C)** Oligodendrocyte counts at 7 dpf in control (gray) and 4 µM late IOX1-treated larvae (0.82 [0.81–0.82] vs 0.83 [0.82–0.84]; p-value = 0.485). **(D)** Oligodendrocyte counts at 7 dpf in control (gray) and 0.5 µM IOX1-treated larvae (0.77 [0.76–0.78] vs 0.78 [0.77–0.79]; p-value = 0.688). **(E)** Oligodendrocyte counts at 7 dpf in control (gray) and 2 µM IOX1-treated larvae (0.76 [0.75–0.77] vs 0.80 [0.79–0.80]; p-value = 0.072). **(F)** Confocal image of 7 dpf *Tg(olig2:dsRed)* larvae treated with DMSO (top) or 4 µM IOX1. Dashed region indicating left optic tectum. Scale bar 100 µm. **(G)** Oligodendrocyte precursor cell densities in the optic tectum at 7 dpf in control (gray) and 4 µM IOX1-treated *Tg(olig2:dsRed)* larvae (2.5 [2.4 – 2.6] vs. 2.2 [2.0 – 2.4] *10^3^ cells/mm^2^; p-value: 0.0087). **(H)** Confocal image of 7 dpf *Tg(isl1a:GFP)* larvae treated with DMSO (top) or 4 µM IOX1. Circled region indicating cranial nuclei nIII and nIV. Scale bar 10 µm. **(I)** Tangential nucleus neuron counts at 7 dpf in control (gray) and 4 µM IOX1-treated *Tg(6*.*7Tru*.*Hcrtr2:GAL4-VP16)*; *Tg(14xUAS:GCaMP6s)*larvae (DMSO vs IOX1: 23 [18 – 25] vs. 27 [20 – 29]; p-value: 0.2222). **(J)** Nucleus of the medial longitudinal fasciculus neuron counts at 7 dpf in control (gray) and 4 µM IOX1-treated *Tg(nefma:hsp70l-LOXP-RFP-LOXP-GAL4)*; *Tg(14xUAS:GCaMP6s)* larvae (DMSO vs IOX1: 4 [3 – 5] vs. 4 [3 – 6]; p-value: 0.8469). **(K)** nIII and nIV neuron counts at 7 dpf in control (gray) and 4 µM IOX1-treated *Tg(isl1a:GFP)* larvae (DMSO vs IOX1: 120 [113 – 127] vs. 139 [121 – 157]; p-value: 0.3095). *indicates p-value < 0.05.

**Figure S2:**
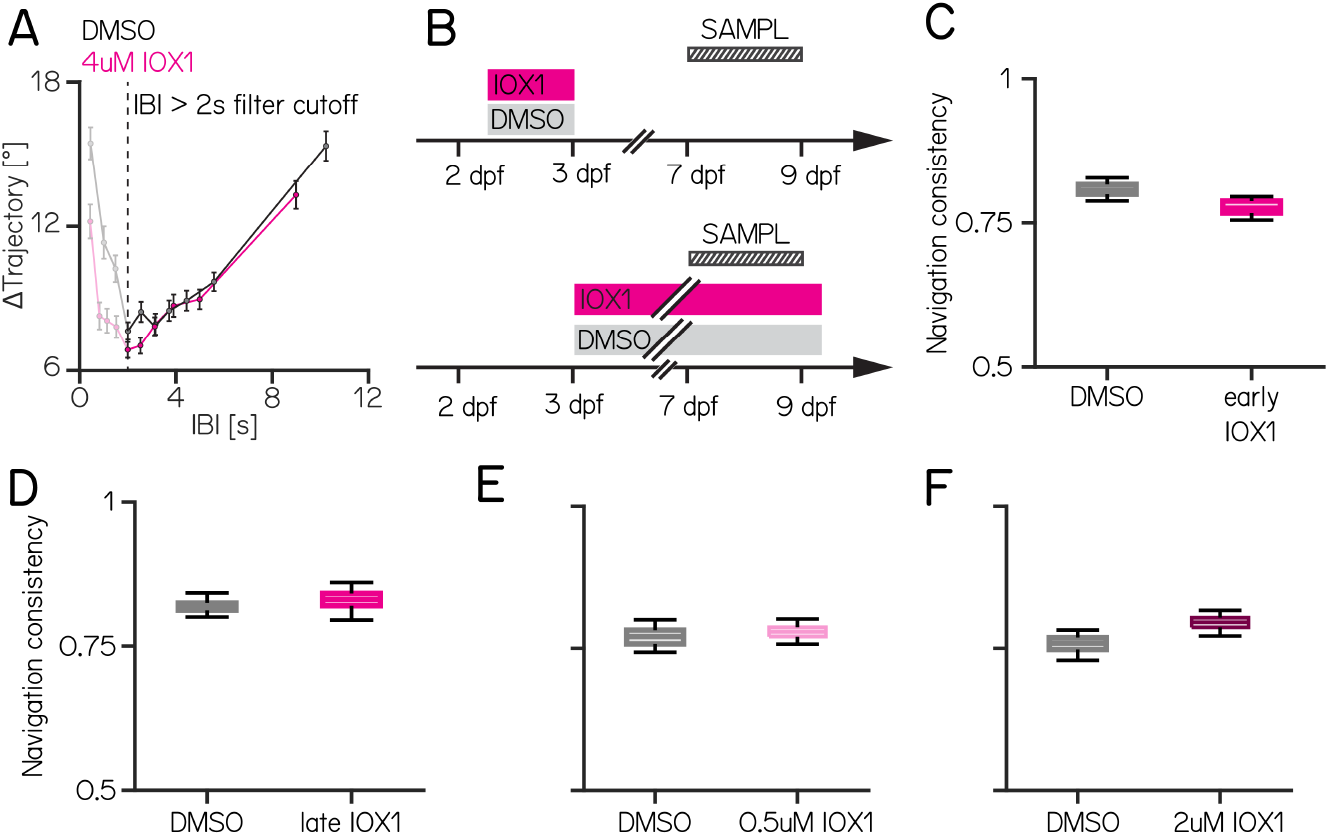
Different timing or concentrations of IOX1 do not affect behavior. **(A)** Change in trajectory (delta) of two consecutive bouts as a function of the inter-bout interval (IBI) illustrating the cutoff used for data analysis. **(B)** Schematic for early (top) and late (bottom) IOX1 treatment. Striped bars indicate the time of behavior testing. **(C)** Navigation consistency for control (gray) and early IOX1 (pink) treated larvae (DMSO vs IOX1: 0.81 [0.80–0.82] vs 0.78 [0.77–0.79]; p-value: 0.228). **(D)** Navigation consistency for control (gray) and late IOX1 (pink) treated larvae (DMSO vs IOX1: 0.82 [0.81–0.82] vs 0.83 [0.82–0.84]; p-value: 0.485). **(E)** Navigation consistency for control (gray) and 0.5 µM IOX1 (light pink) treated larvae (DMSO vs IOX1: 0.77 [0.76–0.78] vs 0.78 [0.77–0.79]; p-value: 0.688). **(F)** Navigation consistency for control (gray) and 2 µM IOX1 (dark pink) treated larvae (DMSO vs IOX1: 0.76 [0.75–0.77] vs 0.80 [0.79–0.80]; p-value: 0.072).

**Figure S3:**
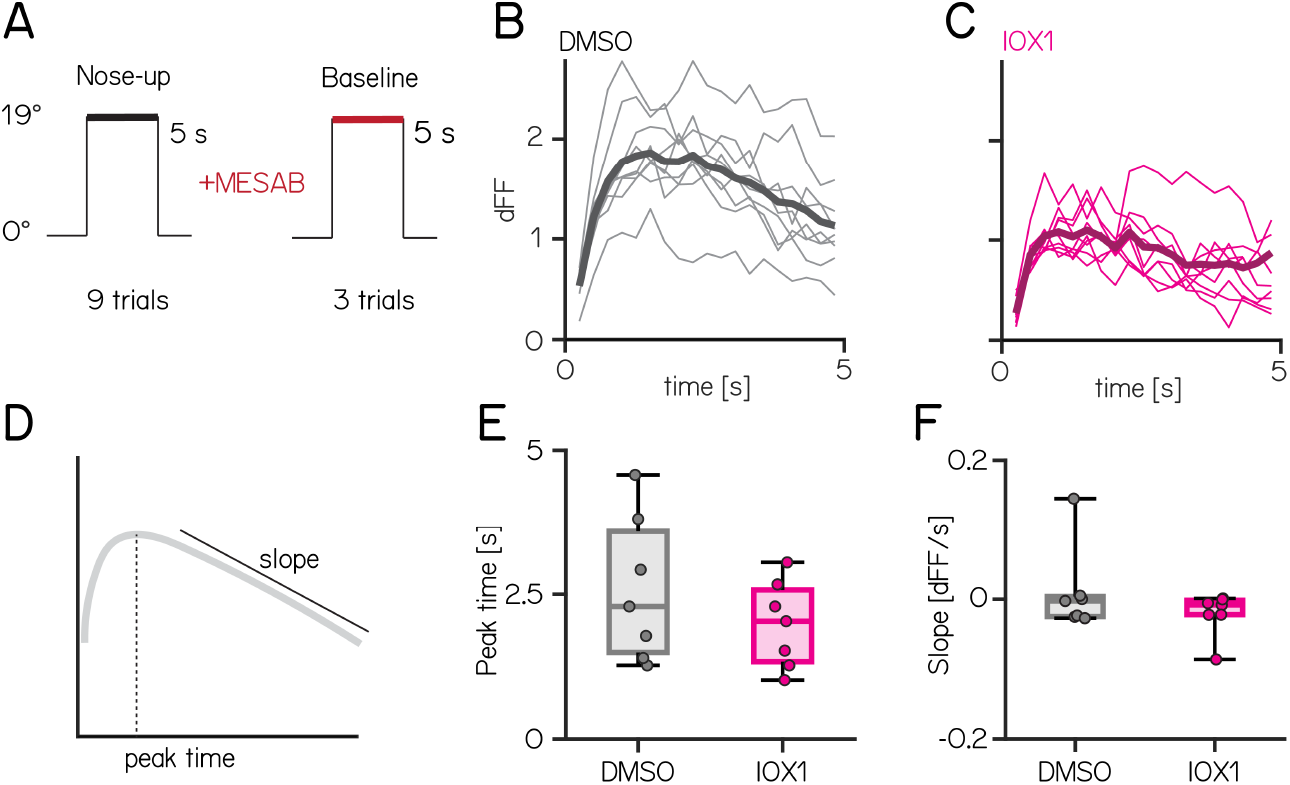
IOX1 treatment does not affect peak fluorescent timing or decay. **(A)** Stimulus trace for eccentric imaging with anesthetized baseline. Cells are imaged during a 5 s stimulus at 19°. To establish a baseline the same cells are imaged in anesthetized animals. **(B)** Example trials of a DMSO treated nMLF/INC neuron, **(C)** Example trials of a IOX1-treated nMLF/INC neuron. **(D)** Schematic of measured values for E - F. **(E)** Quantification of peak time in DMSO versus IOX1-treated animals. (DMSO vs. IOX1: 2.3 [1.5 – 3.6] vs. 2.0 [1.3 – 2.6]; p-value: 0.6591). **(F)** Quantification of GCaMP6s slope in DMSO versus IOX1-treated animals. (DMSO vs. IOX1: −0.00007 [-0.0063 – −0.0010] vs. −0.0021 [-0.0057 – −0.0004]; p-value: 0.7104).

**Figure S4:**
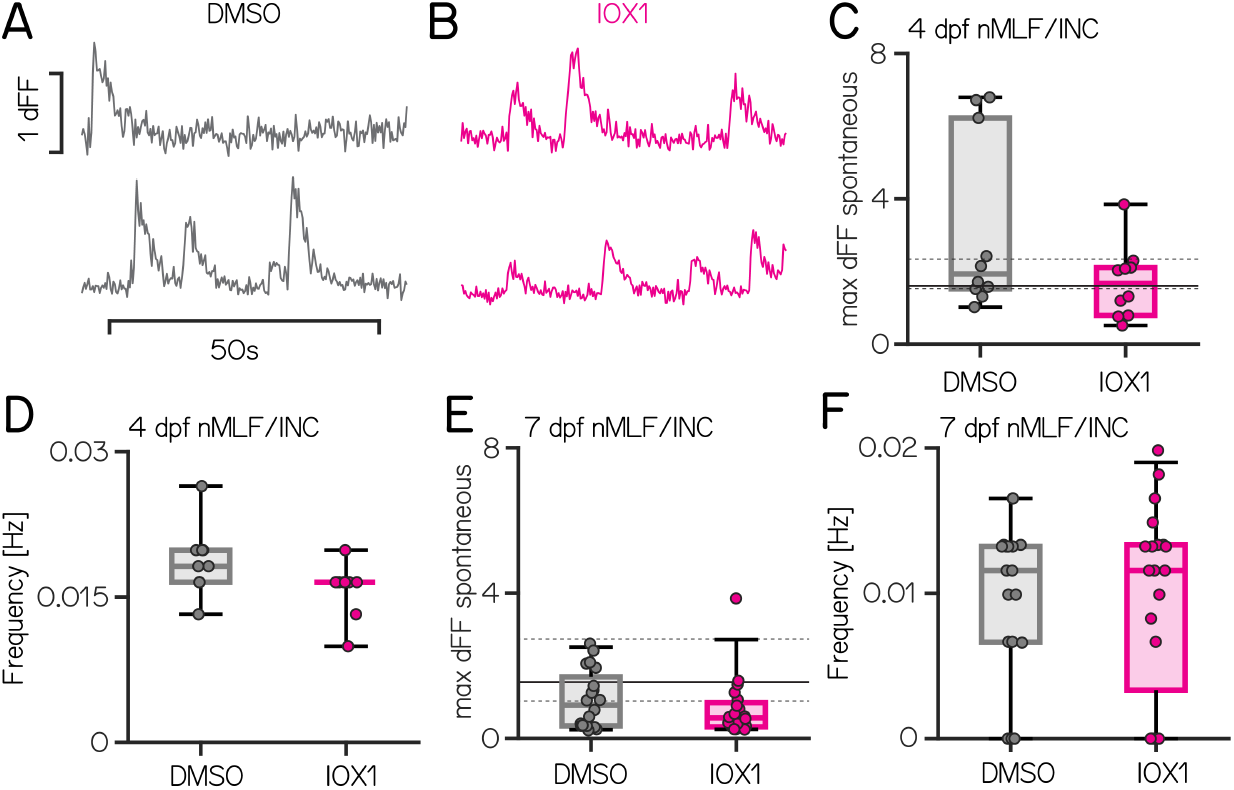
Spontaneous calcium event amplitude or frequency are not affected by IOX1 treatment. **(A–B)** Example traces of spontaneous calcium events in nMLF/INC neurons at 7 dpf in DMSO (gray) or IOX1 treated animals (pink). **(C)** Quantification of spontaneous calcium event amplitude at 4 dpf (DMSO vs. IOX1: 1.93 [1.52 – 6.22] vs 1.67 [0.79 – 2.11]; p-value = 0.162). Full/dashed lines indicate the median and IQR dFF for 4 dpf nMLF/INC nose-up responses. **(D)** Quantification of the frequency of spontaneous calcium events at 4 dpf (DMSO vs IOX1: 0.0182 [0.0165 – 0.0198] vs 0.0165 [0.0165 – 0.0165] Hz; p-value: 0.0514) **(E)** Quantification of spontaneous calcium event amplitude at 7 dpf (DMSO vs. IOX1: 0.92 [0.35 – 1.7] vs 0.57 [0.32 – 0.98]; p-value = 0.3235). Full/dashed lines indicate the median and IQR dFF for 7 dpf nMLF/INC nose-up responses. **(F)** Quantification of the frequency of spontaneous calcium events at 7 dpf (DMSO vs IOX1: 0.0116 [0.0066 – 0.0132] vs 0.0116 [0.0033 – 0.0133] Hz; p-value: 0.7426)

**Figure S5:**
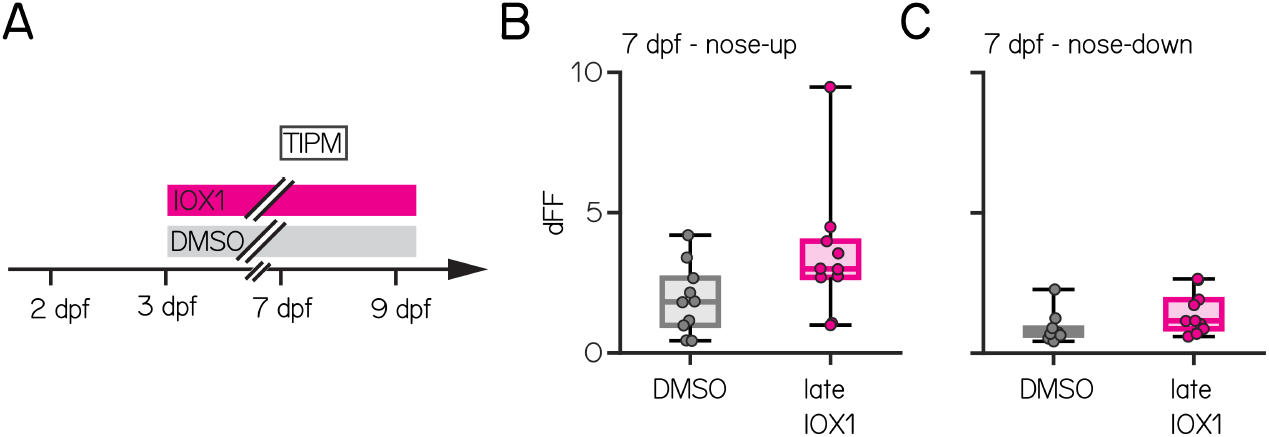
Late IOX1 treatment does not affect nMLF/INC tilt responses. **(A)** Timeline of the experiment showing IOX1 and DMSO-only treatment timecourse beginning at 72 hours post-fertilization. Imaging timepoint is indicated by the open rectangle at 7 days post-fertilization. **(B)** Average responses for 7 dpf nMLF/INC neurons to nose-up stimuli after late IOX1 treatment (DMSO vs late IOX1: 1.82 [0.98 – 2.67] vs. 2.99 [2.69 – 3.98], p-value: 0.0539). **(C)** Average responses for 7 dpf nMLF/INC neurons to nose-down stimuli after late IOX1 treatment (DMSO vs late IOX1: 0.71 [0.64 – 0.89] vs. 1.15 [0.87 – 1.91], p-value; 0.0539).

**Figure S6:**
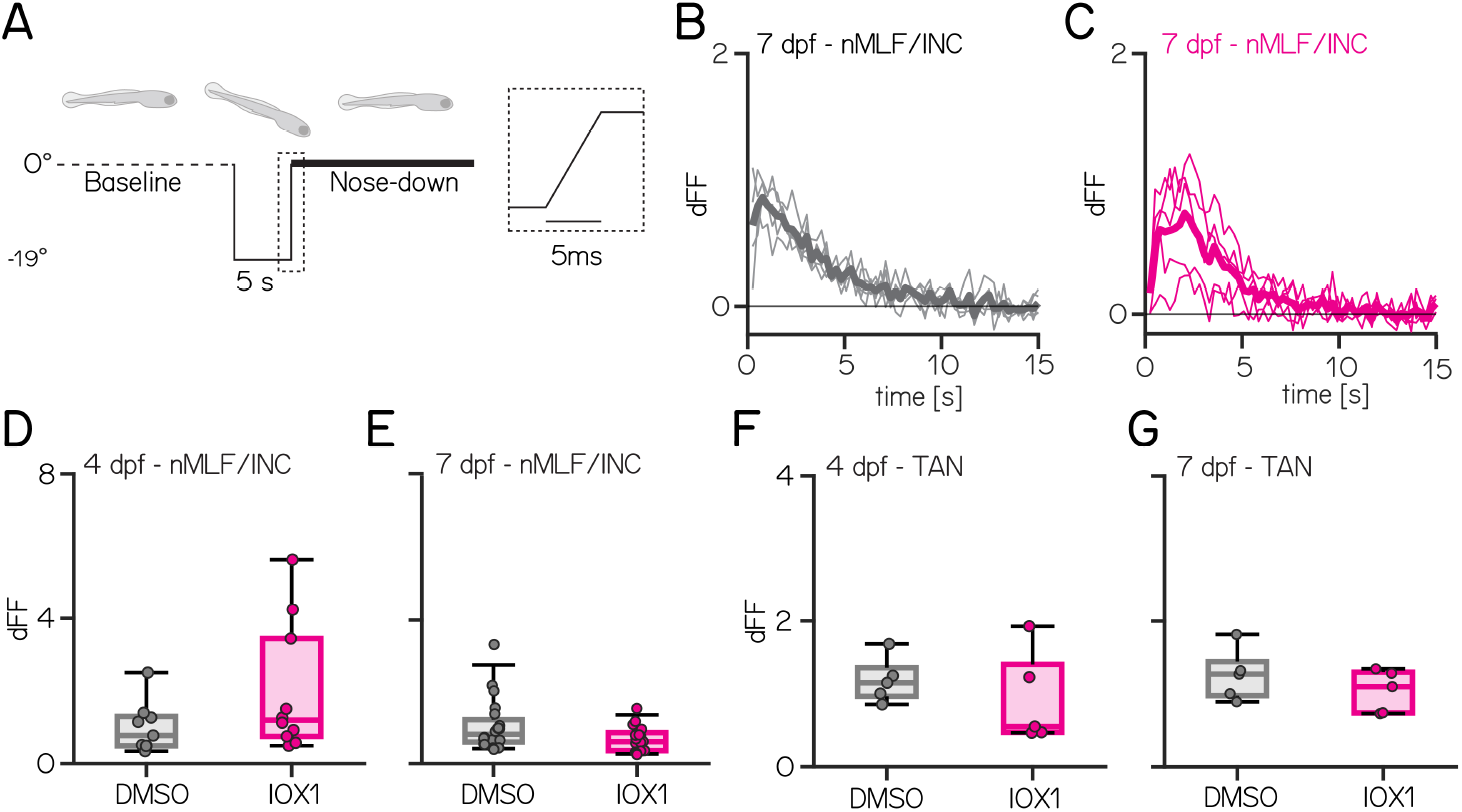
Nose-down pitch tilt responses are not affected by IOX1 treatment. **(A)** Timecourse of nose-down stimulus for imaging indicating pre-tilt baseline (dotted) and post-tilt imaging (thick line). Dotted rectangle corresponds to the −19° return to baseline. **(B-C)** Normalized fluorescent changes after five nose-down tilts in DMSO (E, gray) and IOX1 (F, pink) from 7 dpf fish. Bold line shows the median. **(D)** Average responses for 4 dpf nMLF/INC neurons to nose-down stimuli (DMSO vs IOX1: 0.77 [0.49 – 1.30] vs 1.19 [0.75 – 3.44]; p-value: 0.7124). **(E)** Average responses for 7 dpf nMLF/INC neurons to nose-down stimuli (DMSO vs IOX1: 0.81 [0.60 – 1.22] vs 0.61 [0.36 – 0.86]; p-value: 0.0869). **(F)** Average responses for 4 dpf TAN neurons to nose-down stimuli (DMSO vs IOX1: 1.15 [0.96 – 1.36] vs 0.55 [0.47 – 1.40]; p-value: 0.4206). **(G)** Average responses for 7 dpf TAN neurons to nose-down stimuli (DMSO vs IOX1: 1.27 [0.98 – 1.44] vs 1.10 [0.74 – 1.30]; p-value: 0.5476).

**Figure S7:**
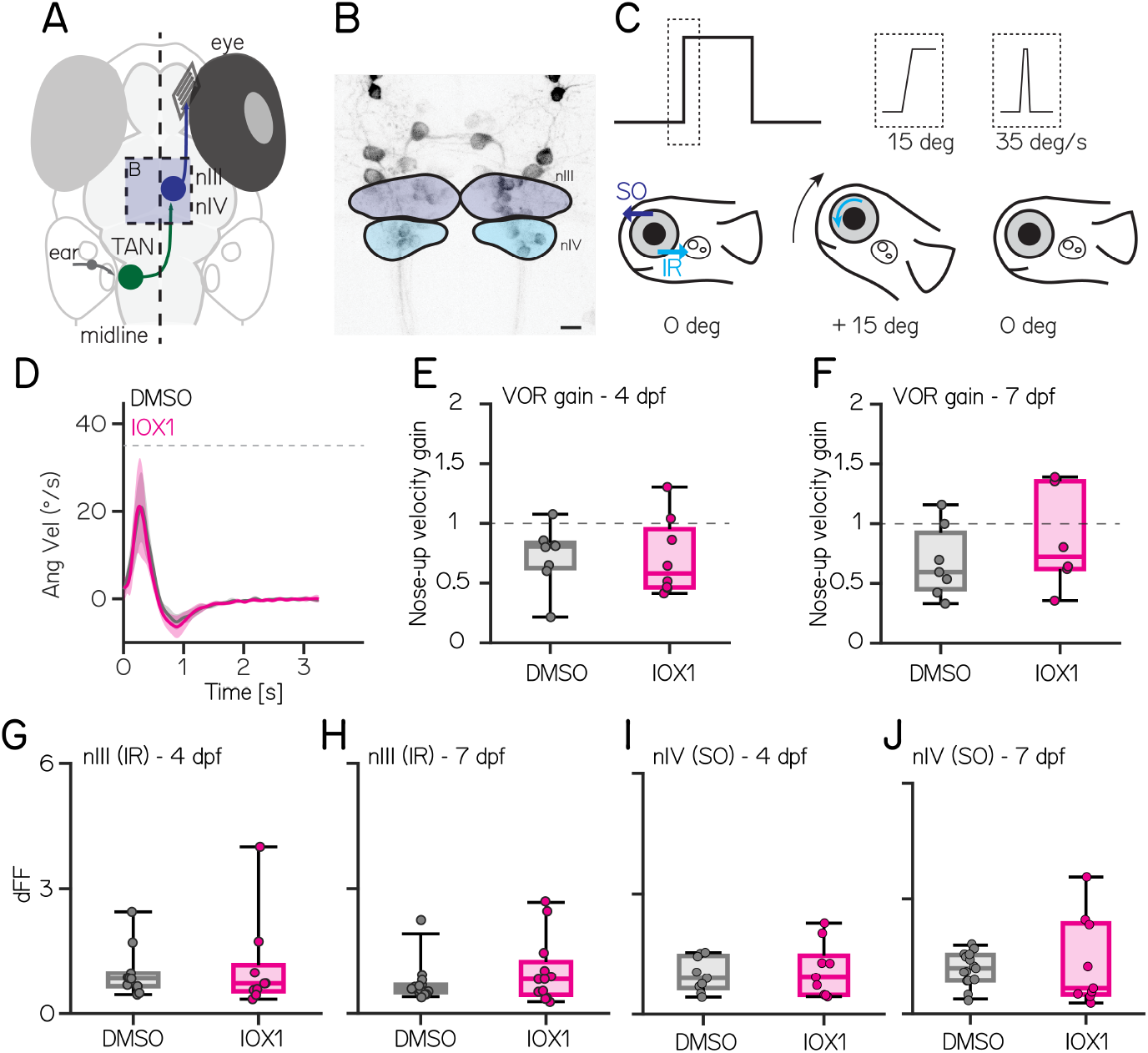
Vestibular ocular reflex circuit and behavior are not affected by IOX1 treatment. **(A)** Schematic of vestibulo-ocular reflex circuit. **(B)** Confocal image of *Tg(nefma:hsp70l-LOXP-RFP-LOXP-GAL4); Tg(UAS:GCaMP6s)* at 7 dpf highlighting the location of nIII (dark blue) and nIV (light blue). Scale bar 10 µm. **(C)** Schematic and details of stimulus for eye movement recordings (top). Fish are rotated to 15° nose-up with a maximum velocity of 35°/s. Fish schematic highlighting the location of the two extra-ocular muscles (superior oblique (SO) and inferior rectus (IR)) that rotate the eyes down in response. **(D)** Average angular velocity traces of 4 dpf DMSO (gray) and IOX1-treated (pink) animals. **(E)** Vestibulo-ocular reflex gain for nose-up stimuli at 4 dpf in DMSO and IOX1-treated larvae (DMSO vs IOX1: 0.81 [0.63 – 0.84] vs 0.58 [0.46 – 0.95]; p-value: 0.7209). **(F)** Vestibulo-ocular reflex gain for nose-up stimuli at 7 dpf in DMSO and IOX1-treated larvae (DMSO vs IOX1: 0.59 [0.45 – 0.92] vs 0.72 [0.62 – 1.35]; p-value: 0.3660). **(G)** Average responses for 4 dpf nIII neurons to nose-up stimuli (DMSO vs IOX1: 0.84 [0.65 – 0.97] vs 0.72 [0.53 – 1.17]; p-value: 0.8809). **(H)** Average responses for 7 dpf nIII neurons to nose-up stimuli (DMSO vs IOX1: 0.59 [0.50 – 0.68] vs 0.82 [0.44 – 1.2]; p-value: 0.4809). **(G)** Average responses for 4 dpf nIV neurons to nose-up stimuli (DMSO vs IOX1: 0.90 [0.65 – 1.46] vs 0.92 [0.48 – 1.47]; p-value: 1.0000). **(H)** Average responses for 7 dpf nIV neurons to nose-up stimuli (DMSO vs IOX1: 1.20 [0.87 – 1.54] vs 0.66 [0.50 – 2.36]; p-value: 0.6764).

